# Interplay between high-energy quenching and state transitions in *Chlamydomonas reinhardtii*: a single-cell approach

**DOI:** 10.1101/2025.01.23.633867

**Authors:** Aliénor Lahlou, Marcelo Orlando, Sandrine Bujaldon, William Gaultier, Eliora Israelievitch, Peter Hanappe, Thomas Le Saux, Ludovic Jullien, David Colliaux, Benjamin Bailleul

**Affiliations:** CPCV, Department of Chemistry, École Normale Supérieure, PSL University, Sorbonne University, CNRS, Paris, France; Paris Research, Sony Computer Science Laboratories, Paris, France; UMR7141: Photobiology and Physiology of Plastids and Microalgae, Institut de Biologie Physico-Chimique, Sorbonne University, CNRS, Paris, France; UMR7144 : Adaptation and Diversity in the Marine Environment (ECOMAP), Station Biologique de Roscoff, Sorbonne University, CNRS, Roscoff, France

**Keywords:** Biophysics of photosynthesis, Single cell physiology, Light stress response in Chlamydomonas reinhartii, Non-photochemical quenching

## Abstract

Studying cell-to-cell heterogeneity is essential to understand how unicellular organisms respond to stresses. We introduce a single-cell analysis framework that enables the study of intercellular heterogeneity of photosynthetic traits, particularly their interactions within individual cells that have identical genotypes, cellular contexts and histories.

Our approach combines single-cell imaging of chlorophyll a fluorescence with machine learning and we study light stress responses in Chlamydomonas reinhardtii as a proof-of- concept.

This framework allows us to score the extent of high-light responses such as state transitions (qT) and high-energy quenching (qE), to reveal significant cell-to-cell heterogeneity and to reveal a strong correlation between qT and qE, undetectable in bulk measurements.

This study highlights the value of single-cell phenotypic analysis for for investigating light stress responses in unicellular organisms. We detail the key aspects that come into play to generalize the method to other complex stress responses involving multiple traits.

## Introduction

It is crucial to study the correlations among traits in biological organisms as they may reveal important trade-offs which arose through evolution between various biological functions. These traits covariations may be difficult to study at the population level because of various factors which are difficult to control like changes in the physiological state of the organisms. Single-cell studies offer significant advantages for understanding the interplay between co-occurring traits in constant genetic background. When trait dispersion is sufficiently high, it provides a continuum of natural trait variations in synchronised isogenic cells in the same environment, a situation that is impossible to achieve when comparing two populations (1).

With regards to single-cell tools, there has been a surge in the development of single-cell sequencing technologies probing “omics” data from genomics, proteomics, metabolomics, lipidomics and glycomics (2), all of them requiring cell lysing. A new challenge is to integrate them with non-invasive physiological measurements and further connect changes in gene expression and proteins, sugar and fat synthesis with physiological characteristics. For this purpose, being able to quantify a physiological process using a non-invasive and specific biological observable is highly desirable.

Chlorophyll a fluorescence (ChlF) emitted by photosystem II (PSII) is common to all photosynthetic organisms. It is the preferred observable for non-invasive studies of photosynthesis as it is highly sensitive and benefits from a century-long history of research (3,4). After light absorption by PSII light-harvesting system, excited chlorophyll can relax through three main de-excitation pathways: photochemistry that initiate photosynthetic electron transfer, heat dissipation or emission of fluorescence – that can be measured optically (5). Their relative efficiencies are determined by the values of the respective rate constants associated to each process. While the rate constant governing fluorescence emission remains invariant, the rate constants of the other processes can be modulated by biological factors. Photochemistry is modulated by the saturation level of the photosynthetic chain, while heat dissipation can be influenced by the regulated production of non-photochemical quenchers in the light-harvesting antenna. Because of such kinetic competition, variations of ChlF can provide valuable information on changes in photochemistry or heat dissipation and is used as a diagnostic tool to assess the overall status of photosynthesis (3,4).

Single-cell photosynthetic responses are also instrumental to improve our basic understanding of photosynthesis. Most of the existing knowledge on photosynthesis has been derived from measurements at the organism or tissue level for plants, and bulk measurements for unicellular organisms (6). Those studies have revealed major insights into the biology of photosynthesis, yet they fail to account for the cell-to-cell heterogeneity, which is observed even within monoclonal populations (7,8). Accessing the biophysical heterogeneity at the single-cell level reveals a richer set of information since the individual responses are not smoothed-out by averaging (1,9–20).

In this work, we selected the response of photosynthetic organisms to high light (HL) stress as a case study. In natural environments, the intensity and wavelength distribution of sunlight fluctuates significantly over time scales ranging from seconds to months (21,22). Several processes mitigate the risk of reactive oxygen species production and PSII photodamage under high light stress, by regulating the photon absorption capacity of the photosystems and/or the efficiency of heat dissipation. These processes induce fluorescence changes and are referred to as non-photochemical quenching (NPQ) (23,24).

We chose to perform the analysis of those processes on the model unicellular green alga *Chlamydomonas reinhardtii* (5) because this organism allows for easy isolation and manipulation of elementary processes involved in high light response. Here, we focus on two NPQ components and their associated biological processes (25):

- High energy quenching (qE), involving the rapidly reversible generation of a non-photochemical quencher in the light-harvesting antenna of PSII and is induced by low luminal pH. In *Chlamydomonas reinhardtii*, qE occurs and relaxes within seconds and is regulated by conformational changes in the Light Harvesting Complexes Stress Related 1 and 3 (LHCSR1 and LHCSR3) induced by low lumenal pH.
- State transitions (qT), involving the reversible movement of a fraction of the light-harvesting complexes between the photosystems. In state I, when all mobile LHCIIs are bound to PSII, the absorption cross-section is large, resulting in high fluorescence levels. In state II, fluorescence decreases when LHCII detaches from to PSII. It occurs and relaxes within minutes, regulated by LHCII phosphorylation via a kinase/phosphatase system (26–29). Although qT is not a form of non-photochemical quenching *per se,* it changes the absorption cross-section of PSII with resulting in variations of ChlF emission (26,27).

We also took into account the contribution of slower NPQ components, in particular photoinhibition (qI), involving photodamaged PSII with a mechanism not yet fully understood (30).

Rather than aiming to resolve the full biological mechanisms behind NPQ regulation, we use the well-characterized responses in *Chlamydomonas* as a testbed to demonstrate how single-cell variability can be used to explore interactions between traits. It is crucial to study the correlations among traits in biological organisms as they may reveal important trade-offs which arose through evolution between various biological functions. Population-level studies typically use reverse genetics (or forward genetic screens) to create knockout mutants of one trait and examine the impact on other traits. However, mutagenesis can lead to secondary mutations and physiological changes, and often fails to capture the complexity of trait interaction, such as nonlinearity or threshold effects. In contrast, single-cell studies offer significant advantages for understanding the interplay between co-occurring traits in constant genetic background. The present study serves as a proof of concept for leveraging the often underestimated benefits of single-cell analyses over population-level studies by exploiting cell-to-cell heterogeneity to gain insights into the interplay between traits.

Our study aims to answer the following questions: *What are the cell-to-cell variations in the extent of a given NPQ component when it is the only active, or between NPQ components when more than one are active? Can we benefit from cell-to-cell variations to explore the interaction between NPQ components?*

Established metrics have been empirically developed to quantify NPQ components from ChlF traces, but they often rely on basic mathematical treatments of specific values such as timestamps, inflection points or local extrema (4,31–34) or more recently on machine learning (35). To better separate contributions from concomitant processes, complementary observables such as fluorescence lifetime or multispectral acquisition can be introduced (36,37) but these strategies come at the cost of complexifying instruments and protocols. In this report, we present a computational method to analyze complex ChlF traces acquired with a CMOS camera and separate the contributions of co-occurring processes. We make the *a priori* strong hypothesis that the behavior of a population (expressing multiple processes) can be satisfactorily accounted for by the referential of populations of a training dataset (expressing one elementary process). By selecting appropriate mutants, choosing specific growth conditions, or using customized treatments, we can build a dataset of reference populations, each exhibiting at most one NPQ component (qE, qT, qI, or none). We trained an algorithm borrowed from the domain of image compression (38) to extract the key ChlF patterns representing elementary kinetics. Then, we introduced an algorithm designed for class-separability problems (39) to project the ChlF traces into a three-dimensional space where each axis represents an NPQ component. This transformation is then applied to wild-type (WT) strains to score the three types of NPQ components (qE, qT, qI). This approach, based on well-characterized responses to light stress, provides clear and unbiased scores for NPQ components in the WT strain. In our case study, we quantify the NPQ components in synchronised isogenic cells to investigate the cell-to-cell heterogeneity of qE and qT separately. Then, we exploit the observed cell-to-cell heterogeneity of qE and qT when they are co-expressed to study the interplay between the underlying biological processes. While this study focuses on qE and qT in *Chlamydomonas*, the framework could be broadly applicable to other species and other types of stress responses involving multiple traits, provided that appropriate training dataset can be generated.

## Materiel and Methods

### Experimental Design: optical calibration

**Microscope set-up:** We developed a fully automated Python-based microscope capable of applying versatile illumination protocols to photosynthetic organisms (leaves or microalgae – see Fig. S1 and Table S1). An Arduino board controls the 470±10 nm and 405±7 nm LEDs at the millisecond resolution. The blue LED is used for the high actinic light, the HL-treatment and the relaxation, and the purple LED is used for the saturating pulses. We chose these wavelengths because it is possible to calibrate precisely their intensity with an actinometer-based protocol (40). A Python algorithm (72,73) segments with a watershed algorithm each cell from the first frame of the movie (Fig. 1A). Each cell mask is used to extract the mean fluorescence of the corresponding alga over the time frames and create a database of *F_m_’* traces for statistical analysis. See Supplementary Information for more details (Fig. S1,S6, Table S1).

**Figure 1:**
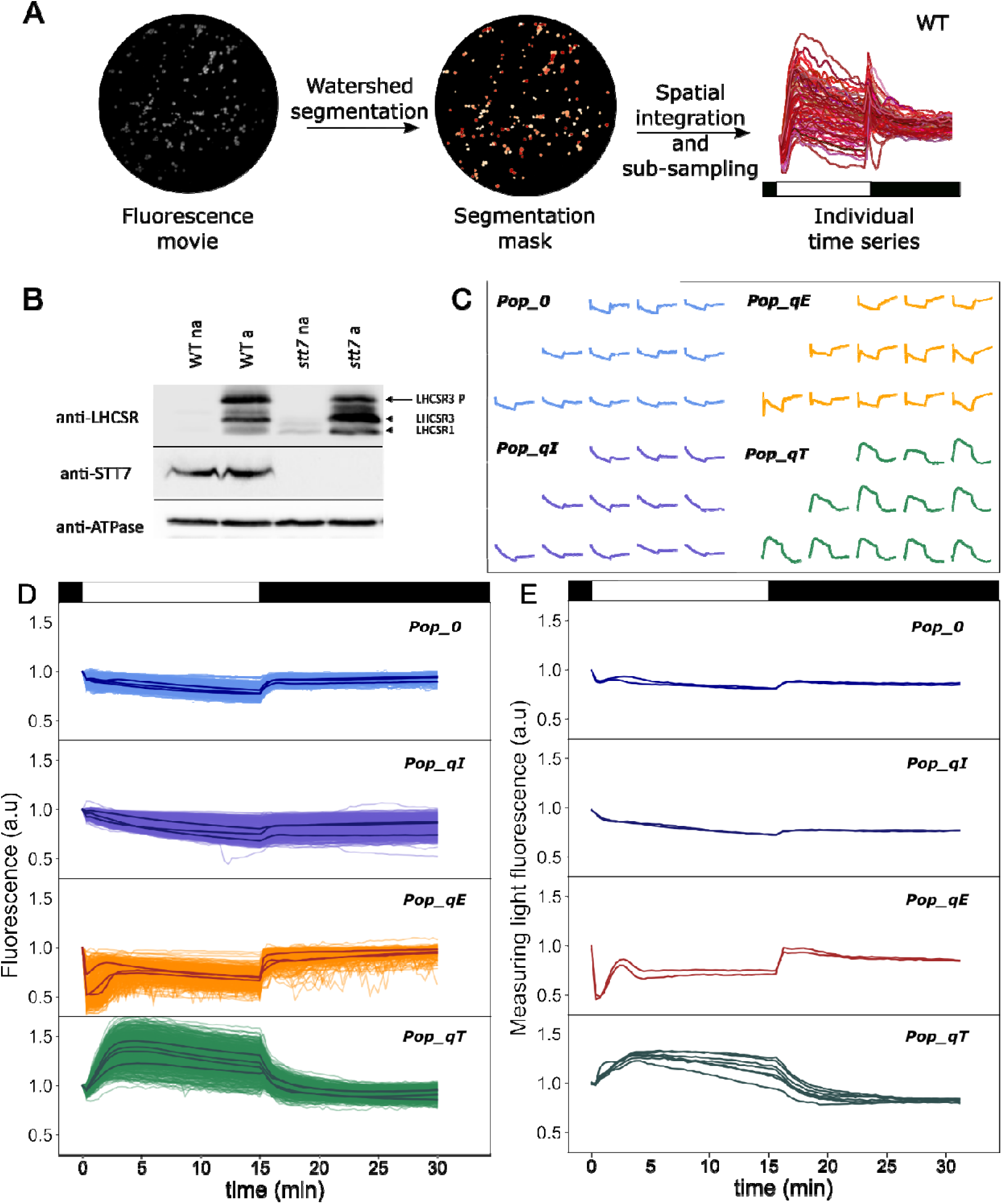
Acquisition of the reference dataset representing elementary ChlF traces for each NPQ component. **A**: *Extraction of the F_m_^’^ traces from single cells of the WT imaged with the fluorescence microscope.* Sample displaying qE and qT: population of the HL-treated WT strain *wt4a^-^*. The reference light protocol (15 min HL-15 min dark) is displayed as a black and white bar. **B**: immuno-quantification of LHCSR (LHCSR1, LHCSR3 and phosphorylated LHCSR3) and STT7 proteins, as well as AtpB (beta- ATP synthase subunit) as a loading control. Wt4a^-^ and STT7 knock-out mutant were sampled as described in Methods, before (na) and after (a) HL-treatment. **C:** Randomly sampled F_m_^’^ traces of the four populations used to build the reference dataset. Samples: populations of ***Pop-0*** (untreated *stt7-1* mutant, 3^rd^ repeat), ***Pop-qI*** (untreated *stt7-1* mutant, 1^st^ repeat), ***Pop-qE*** (HL-treated *stt7-1* mutant, 4^th^ repeat) and (untreated *wt4a^-^*, 4^th^ repeat); **D**: *F_m_^’^ single cell traces of the four populations used to build the reference dataset*. The panels represent the superimposed single cell traces of > 400 cells from at least 3 independent biological replicates (see Table 1 for details) and the average trace is shown as black lines; **E**: *Validation of the microscope setup*. *F_m_’* traces measured on the same four populations as in **D** using a reference macro fluorescence imaging set-up (see Methods) on two independent biological replicates for *stt7*-1 (5 for the *wt4a^-^*). In **D** and **E**, the *F_m_’* data are normalized to the initial *F_m_*, in the dark-adapted cell before the start of the actinic light period. The illumination protocol (HL: white boxes; dark: black boxes) are indicated on the top of the **A**, **D** and **E** panels.

**Table 1:**
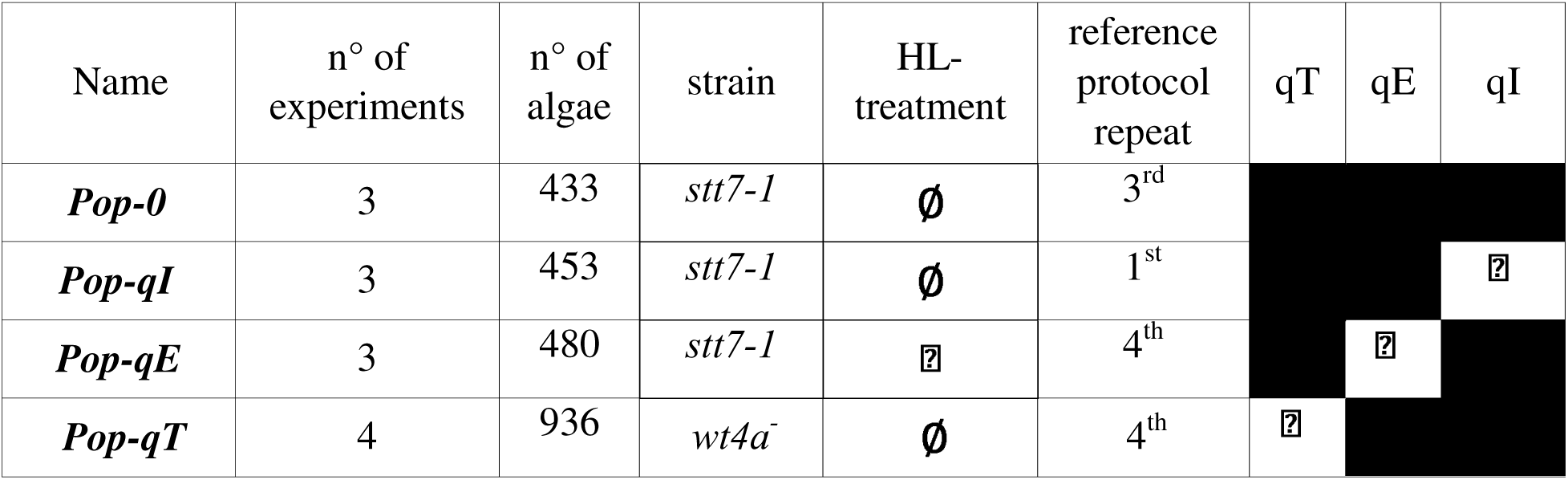
Description of the dataset. (strains, conditioning, and acquisition). ***Pop-0***: untreated *stt7-1* (third repeat of the reference protocol); ***Pop-qI***: *untreated stt7-1* (first repeat); ***Pop-qE****: HL-treated stt7-1* (fourth repeat); ***Pop-qT****: untreated wt4a^-^*(fourth repeat). Shaded cells: NPQ component absent, checkmark: NPQ component present.

**Calibration of the light intensity:** The blue and purple LEDs were calibrated using a point detector (Multi-Pixel Photon Counter). The relaxation time associated to the conversion between ON and OFF state of the Reversibly Photoswitchable Fluorescent Protein Dronpa-2 was monitored (fluorescence filter 525/30), according to the protocol described in (40) and detailed in Fig. S2. Dronpa-2 was extracted and prepared as described in (40): we used 10 *µ*M aqueous solution in Tris buffer pH 7.4 (50 mM Tris, 150 mM NaCl). To obtain the distribution of intensity in the field of view of the microscope, we performed the light calibration of the LEDs by measuring the fluorescence dynamics of Dronpa-2 with a camera (40). The intensity maps and histograms demonstrate the homogeneity of the illumination, with a standard deviation of respectively 15% and 30% of the mean for the 470 and 405 LEDs (see Fig. S2).

**Macro set-up:** Fluorescence emissions were measured using a Speedzen fluorescence imaging setup (JBeamBio, France) at room temperature with actinic light at 620 nm (740 µmol(photons).m^-2^.s^-1^), saturating pulses at 620 nm (5800 µmol(photons).m^-2^.s^-1^, 250 ms) and weak blue “measuring” light pulses (470 nm) to measure the fluorescence yield.

### Experimental Design: Biological samples

**Strains:** The *Chlamydomonas reinhardtii* strains wild type 4A *mt-* (CC-4603), CC-124 wild type *mt*- and *npq4 mt-* (CC-4615) were obtained from the Chlamydomonas Resource Center and *stt7-1 (#a6)* was from the ChlamyStation (41) collection. *stt7-1* is in the same genetic background as *wt4a^-^*. *stt7-1 (#a6)* is a clone allelic to *stt7-1* (42) and non-leaky compared to *stt7-9* (28,43). The strains were synchronized in a 12/12 light cycle (50 µmol(photons).m^-2^.s^-1^ – 23 *°*C) and grown in mixotrophic conditions (Tris-Acetate-Phosphate medium (44)). The synchronized strains grown on Petri dishes were inoculated in liquid medium on Fridays and left to grow under agitation over the weekend. The culture was diluted with fresh Tris-Acetate-Phosphate medium on Mondays and every following days. The experiments were always performed in the exponential phase, between Tuesdays and Fridays. For each experiment, the whole illumination protocol started 2 h after the beginning of the light phase of the photoperiod.

**Sample preparation for microscope experiments:** In all experiments, samples are collected from cultures in exponential phase (from 5*×*10^5^ to 5*×*10^6^ cells/mL), centrifuged (960 g for 8 min) and transferred to Tris-minimal (MIN) medium at approximately 10^7^ cells/mL. The algae were deposited on agarose pads for observations based on protocol from (45). A 1% suspension of agarose in MIN medium was heated to dissolve the agarose. The agarose pads were prepared by dropping 175 µL of melted agarose onto a glass slide holding a spacer *(AB-0578 ThermoFisher)*. To make its surface flat, the pad was covered with a thick microscope glass slide and left for 5 min at 4*°*C for hardening. The cover slide was removed and the preparation was left for 10 min to allow thermal adaptation and evaporation of excess humidity. About 5 µL of algae suspension was dropped onto the agarose and left uncovered for 10 min at room temperature, to remove excess water. Finally, a coverslip was gently placed over the agarose pad for observation. A side opening in the spacer ensured gas exchanges during the experiments (see Fig. S5A).

**Sample preparation for macro set-up experiments:** Cells are harvested in the same conditions as for microscope experiments and go through the same light protocols. They are centrifuged and resuspended in MIN (same protocol as above) at a final concentration of 2*×*10^7^ cells/mL. After at least 30 min of relaxation under the growth conditions they are either used directly or high light pre-treated as explained below. For observations under the SpeedZen, the cell suspensions are placed in a metal plate inside open cavities of 85 µL.

**Monoclonal cultures:** The synchronized monoculture of *stt7-1* and *wt4a**^-^*** were obtained by the streak plate protocol (46). The populations were subcloned once. The mother *wt4a**^-^*** population had not been sub-cloned for more than a year, while the mother *stt7-1* population had been obtained by crossing less than one year before.

**Analysis of polypeptides:** Cells were grown, harvested and conditioned according to the conditions described for macro set-up experiments. The population of *wt4a^-^* and *stt7-1* were harvested (2.10^7^ cells) before (na) and after (a) HL-treatment of four hours. Total cellular proteins were resuspended in 200 mM dithiothreitol (DTT) and 200 mM Na_2_CO_3_ and solubilized in the presence of 2% SDS at 100 °C for 60 s. Cell extracts were loaded on equal chlorophyll basis, separated by SDS-PAGE with 8 M urea (47), and electro-transferred onto nitrocellulose membranes (Amersham Protran 0.1 µm NC) by semi-dry method. Immunodetection was performed using antibodies against LHCSR (48), STT7 (28) and AtpB (beta-ATP synthase subunit) (49). Signals were visualized by enhanced chemiluminescence (ECL - Clarity Biorad) and detected with a luminescent image analyzer ChemidocTM (Biorad).

### Experimental Design: Light protocols

**Reference protocol:** The algae are exposed to 15 min of 400 µmol(photons).m^-2^.s^-1^, 470*±*10 nm, and 15 min of dark. Throughout the whole experiment, saturating pulses (SP) are shone every 20 s (1400 µmol(photons).m^-2^.s^-1^, 405*±*7 nm, 200 ms). The whole fluorescence is collected with a camera, but only the response specific to saturating pulses, *F_m_’*, is analysed. The actinic light is turned off when the SP is applied. The light protocols are illustrated in detail in Fig. S8, S10 and Table S2. The four repeats of the reference protocol and the HL-treatment are performed under the microscope without displacement of the sample. The last two repeats are considered qI-free because the *F_m_* level at the beginning of the exposure is very close to the *F_m_’* level at the end of the exposure (see Fig. S4, S10, S16).

**HL-treatment:** The algae deposited on a pad are exposed to 400 µmol(photons).m^-2^.s^-1^, 470*±*10 nm for 1h20–4 h under the microscope. The HL-treatment is followed by 45 min of 40 µmol(photons).m^-2^.s^-1^, 470*±*10 nm. For macro set-up experiments, the algae suspensions are subjected to 550 µmol(photons).m^-2^.s^-1^ of white light for 4 h under strong agitation. The HL-treatment is followed by 45 min of relaxation under low light (15 µmol(photons).m^-2^.s^-1^, white LED panel).

### Data analysis

Prior to the machine learning protocol, the outliers were removed from the training dataset. The algae with a surface smaller than 5 pixels were discarded. Then, a KD-Tree (50) was built to represent the Euclidean distance relationships between the inputs. We manually set a threshold *D*=0.01 to remove the points with all neighbours further than *D*. In total, 66 fluorescence traces out of 2302 were removed from the training dataset.

**Dictionary Learning** (Fig. 2A, Fig. S11,S12) (38) is used to decompose any complex series of data into pre-learned arbitrary waveforms that capture its most important features. The training phase is unsupervised: it exploits all the *F_m_’* traces from the dataset without any label. It learns to identify characteristic patterns in the ChlF traces that are stored in a dictionary of basic waveforms (called atoms). Its metric is the accuracy of reconstruction of a ChlF trace as a linear combination of the atoms. We used the sparse coding method to decompose the data into a linear combination of atoms from a dictionary (38) implemented in *scikit-learn* (51). The coefficients weighting the atoms form a feature vector. The dimension of this feature vector is lower than the dimension of the original *F_m_’* trace (91) and is equal to *N_D_*, the number of atoms in the dictionary. To build the dictionary, the randomly-sampled equilibrated dataset containing *n_samples_ F_m_’* traces from ***Pop-qE, Pop-qT, Pop-qI*** and ***Pop-0*** was used as a training dataset and fitted with least angle regression following (38). The metric optimized is the reconstruction fidelity with a penalty on the sparsity level, by minimizing the following function:

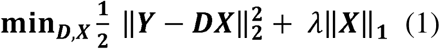

**Figure 2:**
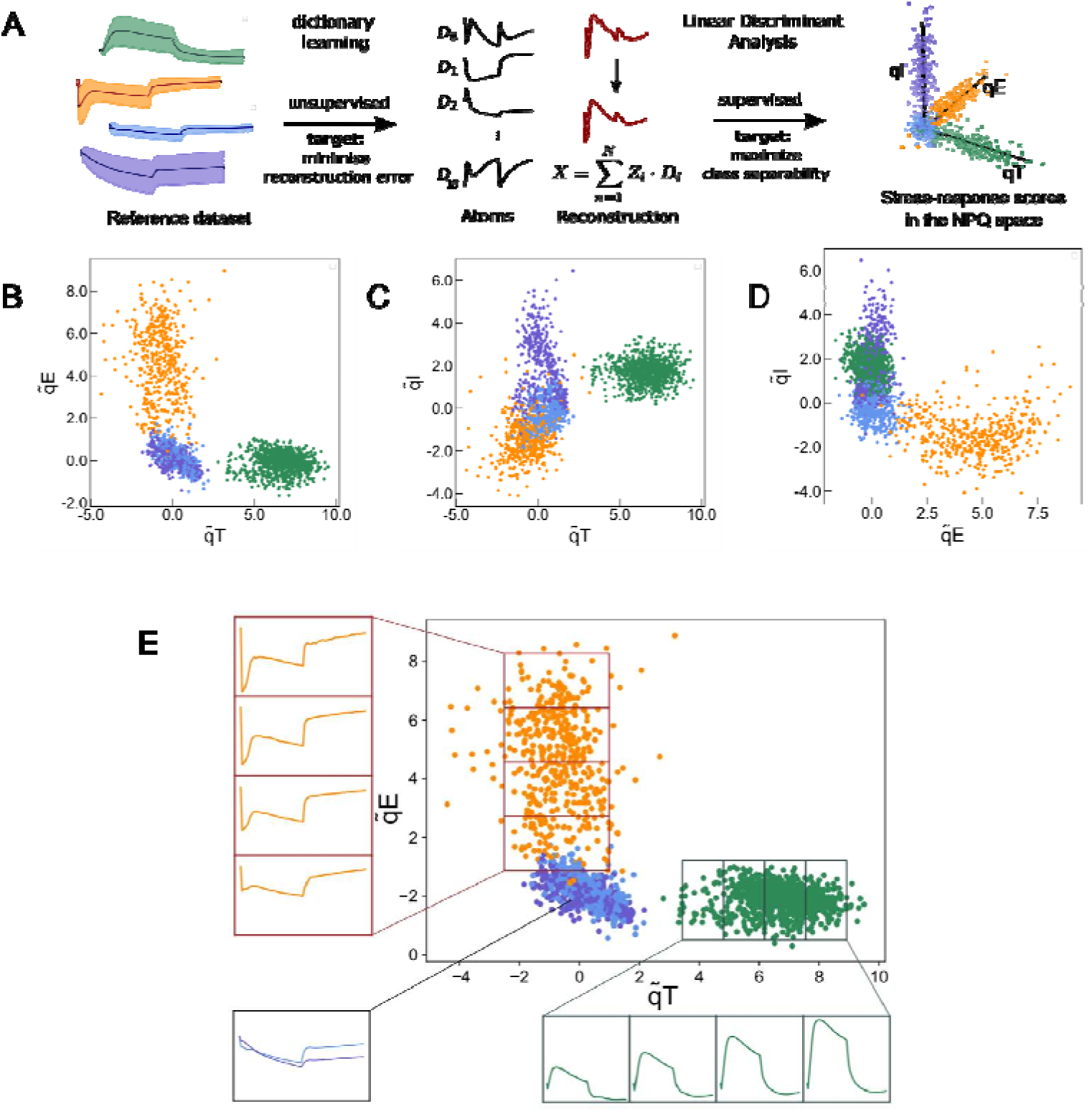
F*r*om *the reference dataset to a base of elementary NPQ components in a 3D space (dataset: see* Table 1). **A**: Flow of the process. A sub-set of the reference dataset traces is first used to perform unsupervised training of an algorithm of dictionary learning which learns to reconstruct the traces with a minimal set of elementary traces (atoms). The resulting reconstructed traces are then combined with the labels of the four training populations to perform a Linear Discriminant Analysis (LDA). It results in a vector in a 3D space, where each axis corresponds to a population of the reference dataset expressing an e ; **B, C, D**: 2D projections of the reference dataset onto the 3D NPQ space after performing Linear Discriminant Analysis on the atom code (dimension 10) from applying dictionary learning. **B** ), **C** . The orange population corresponds to the population ***Pop-qE***. The span of (98% of) the popula 4 equal parts, and the average fluorescence trace within each quarter are computed using the fluorescence traces (normalized to the first point) corresponding to the data points falling inside the quarter. The same is performed for the ***Pop-qT*** . For comparison, the average of the fluorescence traces (normalized to the first point) of the ***Pop-qI*** and ***Pop-0*** populations are also plotted. For all plots, the span of the y-axis is equal to 0.7, with the first point falling at position 1.

Where:

**– Y** is the input data matrix of dimension 91*× n_samples_*,
**– D** is the dictionary matrix with *N_D_* atoms of dimension 91 (normal matrix),
**– X** is the coefficient matrix of dimension *N_D_*,
**–** IIIY- DXII^2^ is the squared Euclidean norm of the reconstruction error,
**–** *||***X***||*_1_ is the L1 norm (sum of absolute values) of the coefficients,
**–** λ is the regularization parameter that controls the sparsity level.

We explored several parameters detailed (Figure S11) including the normalization of the *F_m_’* traces prior to the training. For the analysis described in the Main Text, we selected *N_D_* = 10, *n_samples_*=300, λ = 10*^−^*^6^, and normalized by the sum of the *F_m_’* to obtain a probability distribution.

**Linear Discriminant Analysis** The dimension reduction algorithm, multiclass Linear Discriminant Analysis (39) (LDA) finds the best linear combinations of input features that maximizes the separability among differently-labelled data. The metric optimized is derived from two matrices, *S_B_* and *S_W_* which represent respectively the separation between (B) the means of the classes and the dispersion of the datapoints within (W) each class. We obtain a projection from a space of dimension 10 to a space of dimension *c _−_* 1 where *c* is the number of classes (in our case *c* = 4, ***Pop-qE, Pop-qT, Pop-qI*** and ***Pop-0***). The LDA is solved by Singular Value Decomposition (scikit-learn Python library) (51). Once the projection **T** ∈ 10*×*3 is obtained, we operate a transformation to align the significant axis with the canonical space (*x*, *y*, *z*) for easier analysis. The first step is to identify the principal directions of each class in the dataset and associate it to an axis by collecting the corresponding eigenvector, using Principal Component Analysis (52). They form a transfer matrix **R** which allows to change the base of the space. Then, the origin is identified by using biological knowledge. The average of the point clouds representing ***Pop-qE***, ***Pop-qI*** and ***Pop-0*** is expected to show no qT as the strain *stt7-1* is an STT7 knockout mutant: it is the origin of the q T axis. The average of the point cloud ***Pop-qI*** is expected to show no qE because it has only been exposed to growth light. It is set as the origin of the axis q E. The average of the point clouds ***Pop-qE, Pop-qT*** and ***Pop-0*** are supposed to show no qI since the last *F_m_’* equals the first *F_m_* and are therefore used as the zero reference for the q I axis.

**Evaluation of the cell-to-cell *vs* cell-to-population distances:** (Fig. 3B,E,I) As the experiments are performed on the same sample with *N* algae, we can derive the couple positions in the dictionary learning NPQ space ((*x*_3*,i*_*,y*_3*,i*_*,z*_3*,i*_),(*x*_4*,i*_*,y*_4*,i*_*,z*_4*,i*_))*, i* ∈ [1*,N*] corresponding to algae *i* for the third and fourth repeats. We introduce the distance *D_i,j_* as the distance of the point corresponding to algae *i* in the third repeat to the point corresponding to algae *j* in the fourth repeat:

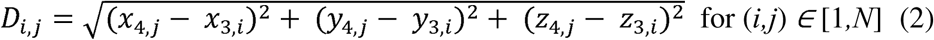

**Figure 3:**
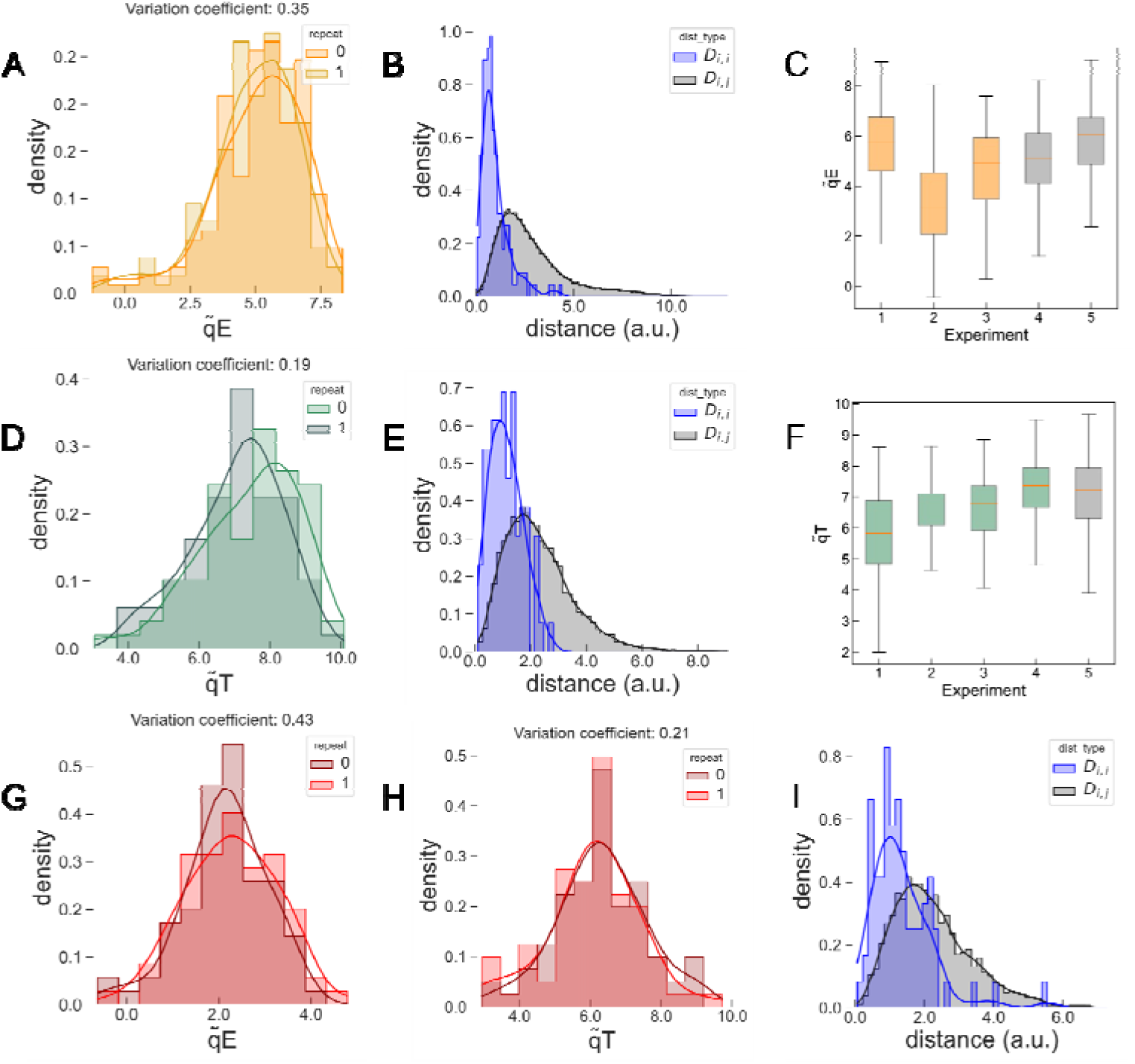
I*n*tercellular *variations in qE and qT in monoclonal synchronised populations*. **A**, **D** (**A,** (**D,** 77 algae) values of two consecutive repeats of the same experiment on isogenic and synchronised cells of the HL-treated *stt7-1* (**A**) and untreated WT *wt4a**^-^*** (**D**) without qI. We evaluate the variation coefficient at 19% for qT, 35% for qE; **B**, **E**: Distributions of the cell-to-cell (*D_i,i_* – blue, SD: 0.7 (**B**), 0.3 (**E**)) *vs* cell-to-population (*D_i,j_* – black, SD: 1.8 (**B**), 1.5 (**E**)) distances in the 3D NPQ space retrieved from **B**: two isogenic HL-treated *stt7-1* populations (2 consecutive repeats; 176 algae); **E**: HL-treated *wt4a**^-^*** population without qI (**E**; 2 consecutive repeats; 77 algae); **C**dataset (orange) and experiments with isogenic synchronized populations (grey); **F** training dataset (green) and experiments with isogenic synchronized populations (grey); **G**, **H** (**G** (**H**) values of two repeats of the same experiment on isogenic and synchronised cells of the HL- treated *wt4a**^-^*** populations without qI. We evaluate the variation coefficient at 21% for qT, 43% for qE; **I**: Distributions of the cell-to-cell (*D_i,i_* – blue, SD: 0.7) vs cell-to-population (*D_i,j_* – black, SD: 1.3) distances in the 3D NPQ space retrieved from two isogenic *wt4a**^-^***populations (2 consecutive repeats; 77 algae). See Table S3 for statistical tests related to the figure.

**Grid representation of the *F_m_* traces in the NPQ space:** To build Fig. 4B,E we grouped the data points in a 8*×*8 grid and plotted the average *F_m_’* trace if there were more than three data points in a grid element. The traces from different point clouds are overlaid with the corresponding colours indicated in Fig. 4A,D.

## Results

### New instrument and protocol to probe high light responses in single cells

We developed a fully automated microscope capable of applying versatile illumination protocols to capture ChlF traces from microalgae immobilized on agarose (see Methods). Our light protocol consists in exposing the algae to high-light and apply saturating pulses (SP) every 20 s to probe the SP-induced fluorescence. Because the yield of photochemistry becomes negligible during SP, the maximal fluorescence yield acts as a reliable indicator of competing non-photochemical processes (25). Since the number of photons exciting the system during the SP is constant, we use the SP-induced fluorescence to track the evolution of the maximal fluorescence yield (*F_m_* before illumination and *F_m_’* during and after HL-treatment, see Methods). We applied an image segmentation algorithm to retrieve the ChlF temporal trace of each alga, called *F_m_’* trace hereafter (Fig S6). In Fig. 1A, we display an example of individual *F_m_’* traces of a wild type (WT) population expressing qE and qT when exposed to our reference protocol which consists in 15 min exposure HL followed by 15 min of relaxation in the dark several times in a row. We chose the strain *wt4a^-^* in this work, otherwise specified.

### Building a reference dataset of elementary qE, qT and qI traces

We developed a machine learning-based workflow to accommodate the variability in the kinetic properties of each NPQ component. Our approach assumes that the ChlF trace of a cell expressing multiple processes, such as in WT (Fig 1A), can be modelled as a linear combination of elementary traces. Therefore, we leveraged extensive knowledge and resources on high light stress responses in *C. reinhardtii* to establish a training dataset, *i.e.* strains and conditions where particular NPQ traits are either present or absent (31,36,53). We defined three distinct populations – specific strains conditioned under appropriate settings – each expressing a single dominant NPQ component, with minimal expression of the others. These populations provided elementary traces representing a single NPQ component, capturing variability in both amplitudes and time constants. Combined with a fourth population lacking all NPQ component, these traces formed the training dataset (see Methods, Table 1, Fig. 1).

For our experiments, we used cells grown under low light in acetate-containing medium to create qE-free conditions (“untreated” hereafter), as these conditions suppress LHCSR gene expression (see western blot in Fig. 1B and (31,32,54)). In contrast, to induce qE, we subjected cells to a 1h20-4h HL-treatment under the microscope after transferring them to acetate-free medium (see Methods), which triggers LHCSR production (“treated” hereafter, Fig. 1B and (31,32,54)). To study conditions without qT, we used the kinase STT7 knockout mutant which remains blocked in state I. We use the WT strain (*wt4a^-^*) to study conditions with qT and the STT7 knockout mutant (stt7-1 #a6, (28)), generated in the *wt4a^-^* genetic background, to study conditions without qT (see STT7 quantification by western blots in Fig 1B).

Based on this, the untreated STT7 knockout mutant was expected to lack both qE and qT. However, we observed a significant fluorescence decrease in this population during the 15 min HL period, which did not relax during the subsequent dark period (see Fig. S10). Several slowly relaxing NPQ components have been described, including qI quenching (30), xanthophyll cycle-dependent qZ quenching (34,55) and sustained qH quenching (56). However, key proteins involved in qH quenching are absent in the *Chlamydomonas reinhardtii* genome (56,57). This slowly induced and slowly relaxing quenching was exacerbated by the chloroplastic translation inhibitor lincomycin (Fig S4), consistent with qI quenching (30). PSII photodamage and qI occur when excessive light disrupts the balance of ongoing degradation and repair processes of reaction centers, particularly the D1 protein encoded in the chloroplast genome (58,59). Therefore, we attribute the slowly reversible quenching observed in our dataset to qI, although a contribution from qZ cannot be strictly excluded (34). Manipulating qI is more challenging than manipulating other NPQ components, as there are no mutant providing qI-free ChlF traces. Although the primary goal of this study was to investigate the heterogeneity and interaction of qE and qT, the inclusion of elementary traces for qI in the training dataset was essential for the machine learning-based approach to work effectively. We found that repeated exposure to our reference protocol progressively reduced the extent of qI, likely reaching a steady-state where PSII repair balanced PSII damage (59). Therefore, for our training dataset, we used the third or fourth iteration of the reference protocol to create conditions without qI contribution and the first iteration to represent conditions with qI.

We used the above considerations to generate the required four populations expressing only qE, only qT, only qI or none of those (Table 1). In practice, we used the 4^th^ repeat of our reference protocol on the untreated WT population to obtain elementary traces of qT (***Pop-qT***), as it expresses the STT7 kinase but not LHCSR3 proteins. The 4^th^ repeat of our reference protocol on the HL-treated *stt7-1* mutant provided elementary traces of qE ***(Pop-qE***) since this mutant then expresses LHCSR3 proteins but lacks STT7 kinase. In the untreated STT7 population, the 1^st^ repeat was used for the elementary traces of qI (***Pop-qI***), whereas the 3^rd^ repeat was used as the reference population displaying none of the NPQ components (***Pop-0***). Indeed, qE slightly increased during the repeats of 15-min illumination needed to reduce the contribution of qI and we came up with using the 3^rd^ repeat instead of the 4^th^ as the best compromise (see Fig 1B, Fig S10b and (31,48,60)).

Fig. 1C displays several randomly sampled traces for the four populations, demonstrating the diversity of the ChlF responses in the training dataset. The whole dataset is plotted in Fig. 1D for ***Pop-qE***, ***Pop-qT***, ***Pop-qI*** and ***Pop-0***. These *F_m_’* traces were collected from movies displaying 100 to 300 algae cells in at least three independent biological replicates (see Table 1, Methods and Fig. S5 for sample preparation details). The algae populations were synchronised but not isogenic (Methods). Mean responses calculated from individual traces of each biological replicate are shown as a black line in Fig. 1D. The data convincingly reproduce the classical trends of the NPQ components (31,32,36,53):

- ***Pop-qE*** traces show a steep decay at the onset of light and rapid relaxation of the fluorescence decline at the end of light consistent with the contribution of qE. In most cells, qE reached a transitory maximum before partially relaxing during the HL period. Such transitory qE has been reported in *Chlamydomonas* (36,48,53) or in plants, where the relaxation of qE in the light was attributed to the concomitant activation of carbon fixation in the Calvin-Benson-Bassham (CBB) cycle (61).
- ***Pop-qT*** traces show a gradual increase of fluorescence at the onset of light and a gradual relaxation during the dark period, both phases lasting a few minutes, indicating qT and the movement of LHCII from PSI (state II) to PSII (state I) in the HL period, and back to PSI in the dark. It might seem counter intuitive that LHCIIs switch from PSI to PSII under high light conditions. Indeed, PSII is already under excessive light pressure and this should favor PQ pool reduction, STT7 activation and state II. However, this behavior has been documented previously
- (31) and attributed to kinase inactivation by negative redox control by the ferredoxin/thioredoxin system (62) or to steric hindrance preventing LHCII kinase phosphorylation due to HL-induced conformational changes in PSII (63).
- ***Pop-qI*** traces show a continuous decline of *F_m_’* during the HL period, which does not fully recover during 15 min of darkness, characteristic of the contribution of qI, with an overall decay between the initial and final *F_m_’*.
- ***Pop-0*** is not completely flat as would be expected if all NPQ components were absent. However, the slight decrease of *F_m_’* at the light onset, representative of qE, and the slight decay of fluorescence between the beginning and the end of the experiment, representative of qI, are both negligible compared to their equivalent in the ***Pop-qE*** and ***Pop-qI***, respectively.

Moreover, these results compare well with the fluorescence traces measured at the population level using a conventional imaging fluorometer (macro set-up, see Methods) by following the same light and dark protocols, including the actinic and saturating pulse (SP) sequences (Fig. 1E) (31,64).

### Using the reference dataset to build a 3-dimensional (qE, qT, qI) space

We used the reference dataset with a machine learning framework to derive a quantitative estimate of the three NPQ components from any ChlF trace. This process involves constructing a 3D space where the three axes represent the elementary biological processes associated with the NPQ components qE, qT, and qI, respectively (3D NPQ space hereafter). The framework performs two consecutive dimension reduction steps, each optimizing a different metric, to transform the *F_m_’* traces into a biologically interpretable 3D vector. Once trained, the framework can be used to analyse new data: an input *F_m_’* trace is first reduced to a vector that describe the linear combination of basic waveforms reconstructing the *F_m_’* trace. This vector is then projected onto a 3D space with axes denoted q T, q E, and q I (Fig. 2A), attributing a score to each NPQ component.

The first step of dimension reduction aims at learning a minimal set of ChlF waveforms from the training dataset to reconstruct generic input ChlF traces with minimal reconstruction error. This step is necessary to validate our working hypothesis stating that a trace presenting co-occurring NPQ components can be interpreted as a combination of elementary traces associated to each NPQ component (see Discussion). We used a dictionary learning method (38) and selected the hyper-parameters to minimise the reconstruction error (see Methods, Fig. S11,12). We selected a reconstruction error threshold (2*_·_*10*^−^*^4^) corresponding to 10 dictionary atoms (Fig. 2A).

The second step is a supervised dimension reduction aimed at maximizing the separability of the annotated training populations. This step learns a matrix to project the output of the first step onto a 3D space using a method based on Linear Discriminant Analysis (LDA, see Methods) (39). LDA learns a projection matrix from the 10-dimensional dictionary output space to a 3D space by identifying three hyperplanes that effectively separate the four training populations. This dimensionality constraint explains the need to select four classes to construct the reference dataset. Once the 3D projection was obtained, we defined the final 3D basis of the NPQ elementary components by assigning the x, y and z axes to the directions along which the training populations ***Pop-qE, Pop-qT*** and ***Pop-qI*** are spread (principal components) and redefining the origin thanks to the population ***Pop-0***. Therefore, the point cloud of each training population extends along a separate axis and, since each population represents an elementary HL stress-response, the axes were named qT , qE components (see Methods).

The 3D projection results of our machine learning framework applied to the training dataset are shown in Fig. 2B,C,D with each panel representing a 2D projection for clarity. The scatterplots of populations expressing the qT or qE component extend only along the qT or qE axis respectively, those not expressing the component overlap at the origin of the corresponding axis. In the plane (q T, q E), the populations ***Pop-qI*** and ***Pop-0*** overlap since they exhibit neither qE nor qT (Fig. 2B). The scatterplot of the population ***Pop-qI*** along the qI axis is also consistent, even though it is less separated from the other populations compared to the other axis (Fig. S16 for details).

The two consecutive steps applied to the training dataset allow the qualitative separation of point clouds corresponding to populations exhibiting distinct NPQ components. Beyond class separation, this representation also allows a quantitative scoring of the NPQ components. Fig. 2E visually demonstrates that the amplitude of the different NPQ components in the fluorescence response increases as the corresponding data point moves away from the origin. For example, the ***Pop-qT*** population extends along the q T axis, with points further from the origin showing a more significant increase in fluorescence at light onset and relaxation to the basal level in the dark. Similarly, the ***Pop-qE*** population exhibits a gradual increase in fluorescence drop at light onset along the q E axis. We present the traces of the ***Pop-qI*** and ***Pop-0*** training populations for comparison and provide the evolution of the fluorescence trace along the q I axis in Fig. S16. We also confirmed that the quantification of the NPQ components based on our machine learning framework matched that based on ad hoc metrics used on bulk populations (31) (see Fig. S13).

### Cell-to-cell variations in qE and qT

We used the 3D projection of traces to assess the cell-to-cell dispersion of ***Pop-qT*** and ***Pop- qE*** populations along the q T and q E axes, respectively. First, we conducted additional experiments to distinguish biological variations from experimental noise (fluorescence noise, instrumental noise, variations of incident light in the field of view, sample heterogeneity or imprecise dictionary reconstruction). While the training dataset consisted in synchronised non-monoclonal populations, we performed new experiments with synchronized monoclonal populations of untreated *wt4a^-^* (***Pop-qT***) and HL-treated *stt7-1* mutant (***Pop-qE***) to minimize genetic diversity and the contribution of cell cycle to intercellular variations (see Methods). To assess noise level, we exploited our microscope setup, which allows us to apply the illumination protocol to the same cell maintained in the same position. We estimated that the distance in the 3D NPQ space between projections of the same cell under two consecutive repeats would provide a good estimate of the experimental noise, assuming the HL stress response remains consistent between the two repeats. Therefore, we compared the distribution of pairwise distance between two consecutive repeats on the same cell (*D_i,i_*) to that between all the other cells in the repeat (*D_i,j_*). If the distance distributions of *D_i,i_* and *D_i,j_* are similar, this suggests that experimental noise governs the scatter of the points cloud. Conversely, if the distribution of *D_i,j_* exceeds that of *D_i,i_* this would indicate that the scatter of the point clouds is mainly due to cell-to-cell biological variations.

First, we observed that the distributions of NPQ scores after two consecutive repeats were statistically similar (Fig. 3A,D), indicating that the average response to HL remained identical (see statistical approach in Fig. S15 and Table S3). We then demonstrated that the distance distribution was twice as narrow for *D_i,i_* as for *D_i,j_* (Fig. 3B,E; a statistical test validated that the distribution variances are different, see Table S3). Thus, we concluded that the scatter of the cloud points is mainly dominated by intercellular biological variations (see Discussion).

The distributions displayed in Fig. 3A,D indicate that the variation coefficients (standard deviation relative to the mean) are ∼40% and ∼20% for qE in ***Pop-qE*** and q T in ***Pop-qT***, respectively. Interestingly, the variation coefficients are not statistically different from those of the non-monoclonal synchronized algae for the reference dataset (Fig. 3C,F), which supports that the distribution is not dominated by a genetic diversity. We further verified that the cell size (estimated from the cell surface in the field of view) did not explain the variance either, and that there was no visible heterogeneity over the sample that could be attributed to the sample design (Fig. S14G,H,I).

### Projection of wild type traces in the 3D NPQ space

We then applied our approach to populations expressing both qE and qT by analysing the projection of single cell *F_m_’* traces from the HL-treated WT *wt4a^-^* in 3D space. First, the monoclonal population *wt4a^-^* was HL-treated for 4 h to induce the expression of qE, in addition to constitutively expressed qT. The average reconstruction error with the dictionary atoms was close to the selected threshold (2.5*_·_*10*^−^*^4^ for 77 cells), validating the hypothesis that the WT response can be accounted for by a combination of the training data. Then, we verified that experimental noise contributed minimally to the dispersion of the point cloud. Fig. 3G,H indicate that successive repeats of the HL protocol on the same cell showed similar mean scores for qE and qT. Moreover, the *D_i,i_* distribution was narrower than *D_i,j_* as previously observed (Fig. 3I), confirming once again that the dispersion is dominated by intercellular biological variations. The mean qE score was lower than that of the untreated *stt7-1* population, consistent with the lower expression of qE in the wild type compared to *stt7-1* mutant as reported in the literature (28,48,65). However, the population exhibited similar coefficients of variation of qE and qT to those observed in training populations expressing only one trait (43% for qE, 21% for qT in Fig. 3G,3H). This significant level of heterogeneity in both NPQ components allowed us to exploit the natural intercellular variations to study the possible interaction between them.

We analyzed how intercellular variations evolve over the qE activation process by applying three sequential HL-treatments of 80 min each, with measurements of *F_m_’* traces after each step, leading to a total of four point clouds in the 3D space. The time evolution of the point clouds in the (q T, qE ) plane is presented in Fig. 4A. Fig. 4B demonstrates that our machine learning framework successively provides quantitative estimates of qE and qT from the complex *F_m_’* traces (Fig. 1A), properly reconstructed by the dictionary method (Fig. S12). Fig 4C shows the evolution of the average (+/- SD) of qE and qT scores, similar to what a bulk measurement would provide. The progressive increase of q E scores with HL exposure time (Fig. 4A,C) aligns with reported results showing that few hours of exposure to HL are needed to reach a steady qE level (31). (Fig. 4A,C). The alignment of affine fits in Fig. 4A demonstrated that qE and qT are strongly correlated, both at the centroids of the scatter plot and among individual cells treated with HL for 80, 160 or 240 min (Fig. 4A), confirmed by statistical tests (Table S3). This observation supports a potential synergistic action of qE and qT in response to HL (5,36,53).

We performed the same experiments with another wild-type strain, *cc124* and found similar results at the population level (Fig. 4D,F). However, unlike *wt4a**^-^***, the correlation observed at the population level (point cloud centroids) is not reflected among individual cells of the same population (Fig. 4D). The *F_m_’* traces displayed in the grid (Fig 4E) were yet properly reconstructed by the dictionary method (Fig S12).

## Discussion

### Comparison to established protocols and quantification of NPQ components

In this work, we used a combination of strains and treatments to generate elementary traces associated with each NPQ component. If the choice of the STT7 mutant to generate traces without qT was obvious, it was more delicate to find appropriate strains and conditions to eliminate qE. To create conditions without qE, we decided to exploit the property that LHCSR proteins are not expressed in a constitutive manner. This choice proved correct since the typical signatures of qE (rapid decrease and increase of *F_m_’* at light onset and offset, respectively) were absent in the ***Pop-qI*** and ***Pop-qT*** populations, which were used to define the qI and qT axis respectively and were not expected to display qE (Fig. 1). Conversely, the qE signature was clearly observed in the *npq4* strain after HL-treatment (Fig. S3), a characteristic previously associated with the presence of the second LHCSR protein, LHCSR1 (54,60). The population ***Pop-0*** is the weak point of our reference dataset, as it presents slight signatures of qE and qI. We made sure our protocol on the new instrument matched results from the literature (Figure S13 and reference (31)).

Our framework requires an initial training phase with a representative population for each phenotype. This limits its application to species for which the necessary resources and expertise are available to build such an informed training dataset. In fact, we experienced this drawback when including the contribution of qI in our analysis, for which the underlying biological processes are not as well controlled and informed as for qE and qT. We see however in (35) that an alternative unsupervised approach can have difficulties to extract generic signatures across wild types and mutants.

One may methodologically question our strong working hypothesis which implies that for every *F_m_’* trace, the kinetics of each NPQ contribution (for example qE) will be reflected somewhere in the collection of single-cell *F_m_’* traces contained in the population representing this NPQ contribution (for example ***Pop-qE*** for qE) in the training dataset. Here, the intermediate reconstruction step with Dictionary Learning assesses the quality of the reconstruction of the *F_m_’* traces by an average mean-squared error test and allows to accept or reject the hypothesis. Failure to reconstruct the traces within the training error margin either invalidates this assumption, or indicates an additional NPQ component not covered by the training dataset. We showed that the *F_m_’* trace of strains exhibiting several NPQ components is effectively captured by a framework built exclusively on populations expressing a single NPQ component (Fig. S12). As shown in Fig. S12, we have confirmed that this hypothesis is applicable to a wide range of strains that were not included in the training dataset and with different genetic background.

We confirmed that the NPQ scores generated by our machine learning framework were consistent with the *ad hoc* measurements commonly used in the literature (Fig. 2,S13,S16), demonstrating the ability of our agnostic method to relate to hand-crafted metrics, offering potential beyond our case study.

### Quality check of the cell-to-cell heterogeneity

We were interested in studying the cell-to-cell heterogeneity of responses to light-stress. We validated our analysis framework to ensure low biases and noise in single-cell responses, preserving our interpretation of intercellular variability or trait correlations through key quality checks that we believe should be reproduced in future works inspired by our approach.

- *Validation of the instrument*. We validated our new instrument and protocols by comparing the results obtained with a classical instrument (Fig. 1D,E). We confirmed the absence of bias related to light heterogeneity or position of the cell in the field of view (Fig. S14).
- *Assessment of the noise impact.* We conducted two successive experiments with each cell assumed to be in identical state, analyzed the cell-to-itself and cell-to-population response variances to evaluate the experimental noise (Fig. 3B,E,I). Achieving identical behavior across all cells in repeated experiments is an unattainable goal, so the noise levels are likely overestimated, which reinforces our conclusions.
- *Reduction of identified sources of biological variation.* Given the reported causal relationships between cell size and photosynthetic activity (66), we confirmed that the NPQ score was not influenced by cell size (Fig. S14D,E,F). Another potential source of variability is related to the cell cycle (67). To minimize this source of variability, we used cultures of cells synchronized on a 12h/12h light/dark cycle (see Methods and Fig. S5,S7). Monoclonal cultures were used when needed to minimize the risk of genetic heterogeneity.

Although we cannot fully exclude other sources of heterogeneity introduced during the cell preparation prior to measurements, we conclude that the observed variability in qE (coefficient of variation ∼0.4) and qT (coefficient of variation ∼0.2) likely reflects stochastic gene expression (68,69).

### Molecular origins of the strong correlation between qE and qT

The substantial intercellular variability of qE and qT allowed to identify a strong correlation between qE and qT in synchronized monoclonal cultures of *wt4a**^-^*** (Fig. 4A). This result is particularly significant because it emerges within the same genotype and cellular context, among cells with common histories. The correlation can be summarized as follows: the higher the qE under HL, the less pronounced the transition from state II to state I.

There are several potential links between qE and qT which allow to identify tentative hypotheses, that are not mutually exclusive and can together explain the strong correlation between qE and qT in the HL-treated *wt4a**^-^***(Fig. 5). First of all, exposure to high light triggers both the induction of the LHCSR3 genes, which drive qE (31,54), and the inactivation of STT7 involved in qT (62,63). The observed anti-correlation between qE and qT at the population level is expected due to HL treatment (Figure 5, “**HL-treatment effect**”). The progressive decline in qT scores over time under HL exposure can be attributed to an increasing proportion of cells locked in state I. However, it does not explain the alignment of qE/qT correlations in the *wt4a^-^* wild-type when comparing the centroids of the scatter plots at different times of HL exposur or all individual cells within a given time point (see Fig 4A). These findings point to a more intricate interaction between qE and qT than can be inferred from bulk measurements, which obscure such details through population-level averaging.

**Figure 4:**
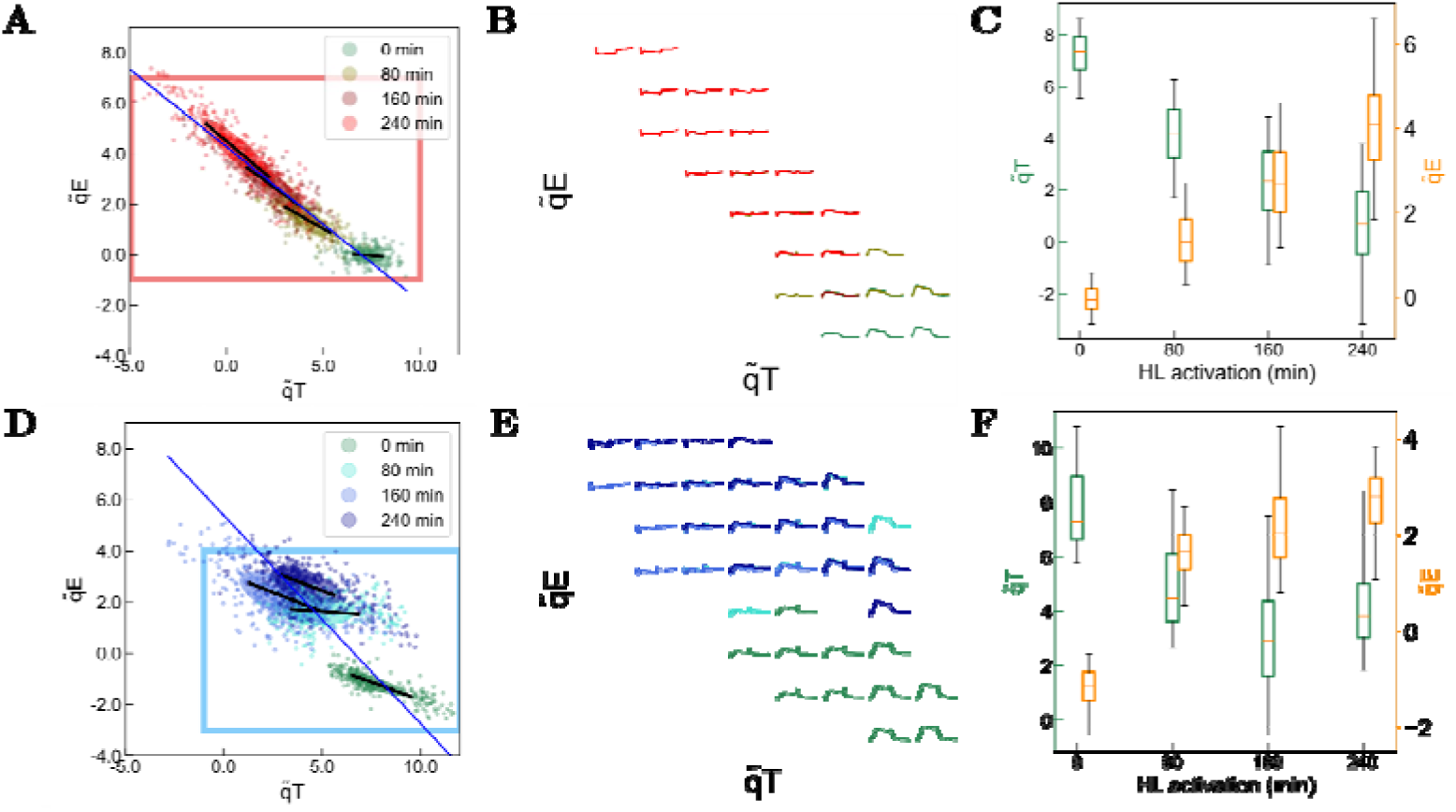
E*v*olution *of the distribution of wild type populations throughout HL-treatment (not isogenic)*. **A**: Wild type *wt4a****^-^*** population taken from the growth light not HL-treated and displaying minimal qI component (untreated *wt4a****^-^*** equivalent to ***Pop-qT***, 4^th^ repeat, 196 cells) exposed to three consecutive HL-treatment protocols of 80 min to promote the expression of LHCSR3 proteins and induce qE. The blue line indicates the direction of the affine fitting curve of the point cloud centroids while the black line indicates the affine curve fit of each point cloud separately; **B**: Grid projection of the *F_m_^’^* traces from **A**; **C**(r ght, orange) throughout the HL-treatment of wild type *wt4a**^-^***. **D**: Wild type *cc124* population taken from the growth light not HL-treated and displaying minimal qI component (4^th^ repeat, 335 cells) exposed to three consecutive HL- treatment protocols of 80 min to promote the expression of LHCSR3 proteins and induce qE. The blue line indicates the direction of the affine fitting curve of the point cloud centroids while the black line indicates the affine curve fit of each point cloud separately; **E**: Grid projection of the *F’_m_* traces from **D**; **F** (right, orange) throughout the HL-treatment of wild type *cc124*. In **B** and **D**, the axis limits have been adjusted to the rectangles (red, A – blue, D) and a trace is represented in the grid only if two or more datapoints fall within the box.

**Figure 5:**
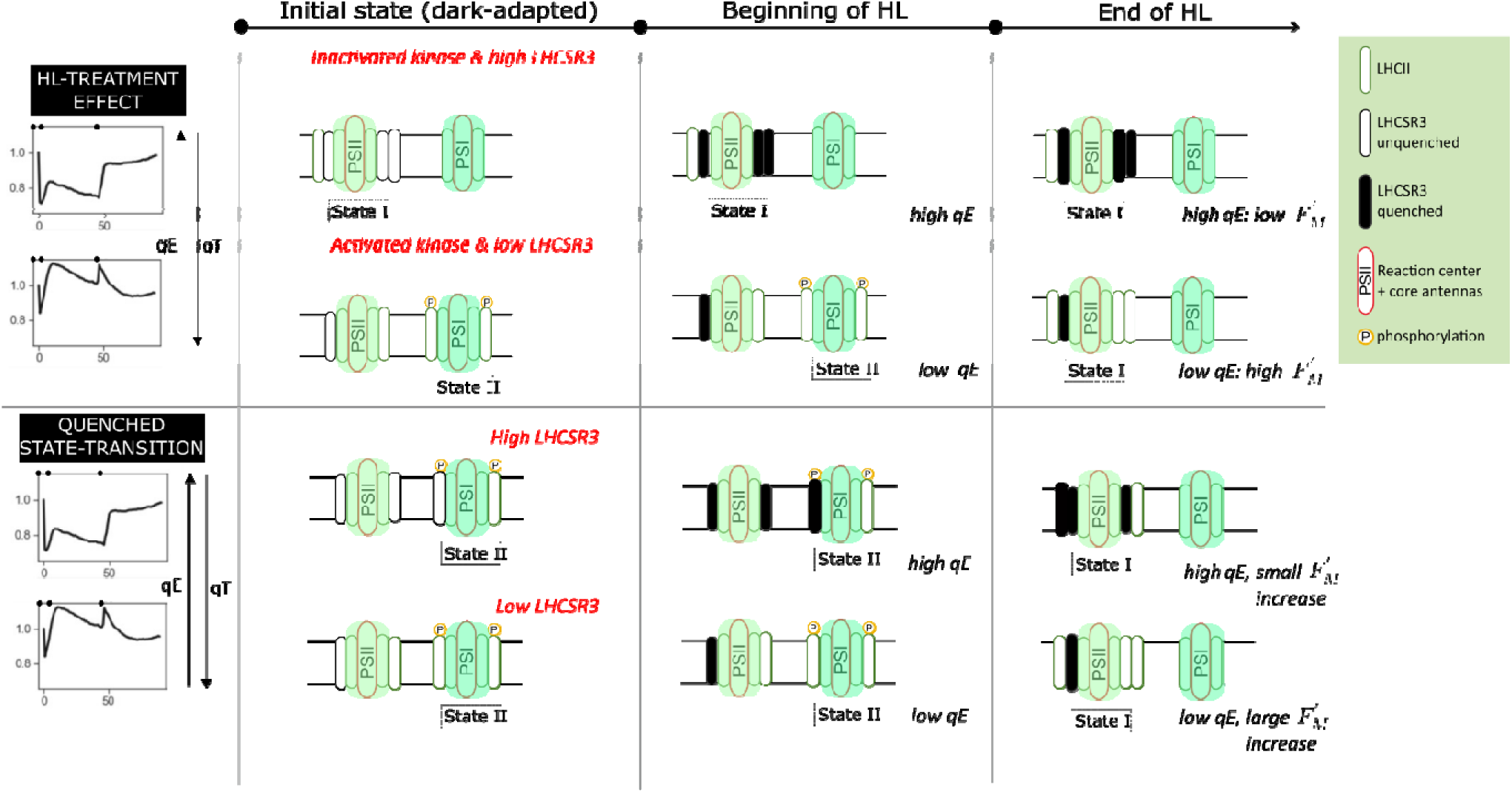
T*w*o *tentative hypotheses accounting for the observed correlation between qE and qT.* The “initial state (dark-adapted)” column represents the initial state of the dark-adapted sample (first *F_m_’* at the beginning of the light protocol). The “beginning of HL” column represents the situation after less than 1 min of HL (first *F_m_’* after the onset of HL). The “End of HL” column represents the situation at the end of the HL period (last *F_m_’* before offset of light).

Although exploring the molecular basis of this correlation is outside the scope of this study, we would like to propose a few tentative hypotheses. One possibility is that the high light signaling pathways, which respectively lead to reduced STT7 activity and to increased LHCSR3 amounts, are coupled. In this scenario, cells with higher LHCSR3 would also exhibit a more pronounced reduction in the accumulation or activity of STT7. Alternatively, a mechanistic coupling between qE and qT could exist, where, for the same amounts of LHCSR3 or STT7, the extent of state transitions influences heat dissipation in PSII, or vice versa. An obvious mechanistic coupling between the two processes arises from the fact that, for a given amount of LHCII transferred from PSI to PSII, the resulting fluorescence increase will be smaller (lower qT score) if PSII fluorescence is strongly quenched (higher qE score). In this "**quenched state-transitions**" hypothesis, an initial cell-to-cell heterogeneity in the amount of LHCSR3 bound to PSII (reflecting variability in qE capacity) leads to a corresponding variation in fluorescence increase for each LHCII complex moving from PSI to PSII. The reported phosphorylation of LHCSR proteins by STT7 could also participate to this coupling (28,31,48). Other potential mechanistic links may involve Cyclic Electron Flow (CEF) around PSI or the activation state of the CBB cycle.

Regardless of the molecular causes for the different behaviors of the *wt4a^-^* and *cc124* wild-types, their comparison provides two insights. First, it highlights the value of single cell studies, as two populations may orchestrate their responses to high light differently, while displaying similar behaviour at the macro (population) level. Second, it highlights the potential influence of the Yule-Simpson paradox in photosynthesis research, where a correlation observed at the population level may not reflect individual causal relationship (70).

### Perspectives for single-cell photosynthesis measurements

We demonstrated that our machine learning framework is able to predict stress-response levels without a priori knowledge of the temporal phases, but taking into account only the dynamics of the signal. As the framework is agnostic to the biological question, it can be readily extended to other single- cell phenotyping applications involving kinetic responses, across different species and stress types. For instance, it could be adapted to study responses to nutrient deficiencies or extreme temperatures, provided that these stresses trigger detectable changes in fluorescence yields.

Importantly, this single-cell machine learning approach allows to derive information on stress response heterogeneity within populations with a small number of experiments. It also allows for the identification of statistically significant correlations between different stress responses, highlighting an underlying biological interplay. One promising application is the study of the interplay between PsbS- and LHCSR3-dependent NPQ components in mosses. Moreover, the same strategy could be used to explore interactions between photochemical quenching components, rather than focusing only on non-photochemical quenching. Provided the method is sensitive enough under lower light intensities than those typically used to probe NPQ, it could, for example, be applied to investigate the interaction between CEF and the flavor-diiron protein pathway in *C. reinhardtii*.

Future work could integrate ChlF measurements with flow cytometry/microfluidics and/or omics methods to correlate cell morphology and photosynthetic traits, facilitating applications like varietal selection, directed evolution, and single-cell screening.

## Acknowledgments

The authors would like to thank Francis-Andre Wollman, Olivier Vallon, Stephan Eberhard, Julien Sellés, Pierre Neveu, Vincent Croquette and David Bensimon for helpful discussions, Marie-Aude Plamont for preparing Dronpa-2 extracts, Douglas Boari for participating in the Arduino script development, and Sébastien Marino for illustrations.

## Author contributions

Conceptualization: AL, DC, LJ, TLS, BB Methodology: AL, MO, SB, PH, DC, LJ, TLS, BB

Software: AL, WG, PH, DC Validation: AL, MO, SB, DC, BB Formal analysis: AL, MO, SB, DC, BB

Investigation: AL, MO, SB, EI, PH, DC, LJ, TLS, BB Resources: AL, MO, SB, EI, PH, DC, LJ, TLS, BB

Data curation: AL, MO, WG, DC Writing – original draft: AL, LJ, DC, BB

Writing – Review and editing: AL, MO, SB, WG, EI, PH, DC, LJ, TLS, BB Visualisation: AL, MO, SB, WG, DC, BB

Supervision: AL, PH, DC, LJ, TLS, BB Project Administration: LJ, TLS, PH, DC, BB Funding acquisition: LJ, TLS, PH, DC, BB

## Data and materials availability

All data are available in the main text, the supplementary materials or the Github repository https://github.com/DreamRepo/NPQScore-data. DOI: https://doi.org/10.5281/zenodo.14606011

SI references: (31,32,34,40,45,53,54,58–60,71–77)

## Abbreviations

ChlF: endogenous chlorophyll a fluorescence
PSII: photosystem II
NPQ: Non-Photochemical Quenching of chlorophyll fluorescence
LEF: Linear Electron Flow
CEF: Cyclic Electron Flow
CBB cycle: Calvin-Benson-Bassham cycle
MIN: Tris-minimal culture medium
WT: wild type
STT7: state-transition 7 kinase
LHCII: Light Harvesting Complex II
LHCSR: Light Harvesting Complex Stress Related
PAM: Pulse Amplitude Modulated
SP: Saturating Pulse of light
HL: High Light
Fm’: maximal fluorescence yield in light-adapted sample
Fm: maximal fluorescence yield in dark-adapted sample
qT: NPQ component due to state-transition
qE: energy-dependent quenching
qI: photoinhibition-related quenching
Pop-0: population of stt7-1 without HL-treatment without qI
Pop-qE: population of stt7-1 with HL-treatment without qI
Pop-qI: population of stt7-1 without HL-treatment with qI
Pop-qT: population of wild type wt4a- without HL-treatment without
qI LDA: Linear Discriminant Analysis
3D NPQ space: three-dimensional space with axis defined by elementary NPQ components
qT: axis of the 3D NPQ space defined by Pop-qT
qE: axis of the 3D NPQ space defined by
Pop-qE qI: axis of the 3D NPQ space defined by Pop-qI

## References

1. Damodaran SP, Eberhard S, Boitard L, Rodriguez JG, Wang Y, Bremond N, et al. A millifluidic study of cell-to-cell heterogeneity in growth-rate and cell-division capability in populations of isogenic cells of Chlamydomonas reinhardtii. PloS one. 2015;10(3):e0118987.

2. Miao Z, Humphreys BD, McMahon AP, Kim J. Multi-omics integration in the age of million single-cell data. Nature Reviews Nephrology. 2021 Nov;17(11):710–24.

3. Krause GH, Weis E. Chlorophyll fluorescence as a tool in plant physiology: II. Interpretation of fluorescence signals. Photosynthesis Research. 1984;5(2):139–57.

4. Papageorgiou GC, Govindjee, Govindjee, editors. Chlorophyll a Fluorescence: A Signature of Photosynthesis. Dordrecht: Springer Netherlands; 2004. (Advances in Photosynthesis and Respiration; vol. 19).

5. Grossmann AR, Wollman FA, editors. The chlamydomonas sourcebook. Volume 2: Organellar and metabolic processes. Third edition. London, UK San Diego, CA Cambridge, MA Oxford, UK: Academic Press, An imprint of Elsevier; 2023.

6. Cole B, Bergmann D, Blaby-Haas CE, Blaby IK, Bouchard KE, Brady SM, et al. Plant single-cell solutions for energy and the environment. Communications biology. 2021;4(1):1–12.

7. Brehm-Stecher BF, Johnson EA. Single-cell microbiology: Tools, technologies, and applications. Microbiology and molecular biology reviews. 2004;68(3):538–59.

8. Lidstrom ME, Konopka MC. The role of physiological heterogeneity in microbial population behavior. Nat Chem Biol [Internet]. 2010 Oct [cited 2023 Jun 30];6(10):705–12. Available from: https://www.nature.com/articles/nchembio.436

9. Snel JFH, Dassen HHA. Measurement of Light and pH Dependence of Single-Cell Photosynthesis by Fluorescence Microscopy. Journal of Fluorescence. 2000 Sep;10(3):269–269.

10. Konert G, Steinbach G, Canonico M, Kaňa R. Protein arrangement factor: A new photosynthetic parameter characterizing the organization of thylakoid membrane proteins. Physiologia plantarum. 2019;166(1):264–77.

11. Westerwalbesloh C, Brehl C, Weber S, Probst C, Widzgowski J, Grünberger A, et al. A microfluidic photobioreactor for simultaneous observation and cultivation of single microalgal cells or cell aggregates. PLOS ONE. 2019 Apr;14(4):e0216093.

12. Behrendt L, Salek MM, Trampe EL, Fernandez VI, Lee KS, Kühl M, et al. PhenoChip: A single-cell phenomic platform for high-throughput photophysiological analyses of microalgae. Science advances. 2020;6(36):eabb2754.

13. Andersson M, Johansson S, Bergman H, Xiao L, Behrendt L, Tenje M. A microscopy-compatible temperature regulation system for single-cell phenotype Analysis–Demonstrated by thermoresponse mapping of microalgae. Lab on a Chip. 2021;21(9):1694–705.

14. Herdean A, Sutherland DL, Ralph PJ. Phenoplate: An innovative method for assessing interacting effects of temperature and light on non-photochemical quenching in microalgae under chemical stress. New Biotechnology. 2022;66:89–96.

15. Küpper H, Šetlík I, Trtílek M, Nedbal L. A microscope for two-dimensional measurements of in vivo chlorophyll fluorescence kinetics using pulsed measuring radiation, continuous actinic radiation, and saturating flashes. Photosynthetica. 2000;38(4):553–70.

16. Šetlíková E, Šetlík I, Küpper H, Kasalickỳ V, Prášil O. The photosynthesis of individual algal cells during the cell cycle of Scenedesmus quadricauda studied by chlorophyll fluorescence kinetic microscopy. Photosynthesis research. 2005;84(1):113–20.

17. Trampe E, Kolbowski J, Schreiber U, Kühl M. Rapid assessment of different oxygenic phototrophs and single-cell photosynthesis with multicolour variable chlorophyll fluorescence imaging. Marine Biology. 2011 Jul;158:1667–75.

18. Mohr W, Vagner T, Kuypers MM, Ackermann M, LaRoche J. Resolution of conflicting signals at the single-cell level in the regulation of cyanobacterial photosynthesis and nitrogen fixation. PLOS one. 2013;8(6):e66060.

19. Gachon CMM, Küpper H, Küpper FC, Šetlík I. Single-cell chlorophyll fluorescence kinetic microscopy of Pylaiella littoralis (Phaeophyceae) infected by Chytridium polysiphoniae (Chytridiomycota). European Journal of Phycology. 2006 Nov;41(4):395–403.

20. Széles E, Nagy K, Ábrahám Á, Kovács S, Podmaniczki A, Nagy V, et al. Microfluidic Platforms Designed for Morphological and Photosynthetic Investigations of Chlamydomonas reinhardtii on a Single-Cell Level. Cells. 2022 Jan;11(2):285.

21. Pearcy RW. Sunflecks and photosynthesis in plant canopies. Annual review of plant biology. 1990;41(1):421– 53.

22. Murchie EH, Kefauver S, Araus JL, Muller O, Rascher U, Flood PJ, et al. Measuring the dynamic photosynthome. Annals of botany. 2018;122(2):207–20.

23. Minagawa J, Tokutsu R. Dynamic regulation of photosynthesis in Chlamydomonas reinhardtii. The Plant Journal. 2015;82(3):413–28.

24. Ruban AV. Nonphotochemical chlorophyll fluorescence quenching: Mechanism and effectiveness in protecting plants from photodamage. Plant physiology. 2016;170(4):1903–16.

25. Goss R, Lepetit B. Biodiversity of NPQ. Journal of Plant Physiology. 2015 Jan 1;172:13–32. Available from: https://www.sciencedirect.com/science/article/pii/S0176161714000686

26. Wollman FA. State transitions reveal the dynamics and fexibility of the photosynthetic apparatus. The EMBO Journal. 2001 Jul;20(14):3623–30.

27. Goldschmidt-Clermont M. Chapter 24 - State transitions. In: Grossman AR, Wollman FA, editors. The Chlamydomonas Sourcebook (Third Edition). London: Academic Press; 2023. p. 787–805. Available from: https://www.sciencedirect.com/science/article/pii/B9780128214305000055

28. Bergner SV, Scholz M, Trompelt K, Barth J, Gäbelein P, Steinbeck J, et al. STT7-Dependent Phosphorylation Is Modulated by Changing Environmental Conditions, and Its Absence Triggers Remodeling of Photosynthetic Protein Complexes. Plant Physiology. 2015 Jun;168(2):615–34.

29. Cariti F, Chazaux M, Lefebvre-Legendre L, Longoni P, Ghysels B, Johnson X, et al. Regulation of Light Harvesting in Chlamydomonas reinhardtii Two Protein Phosphatases Are Involved in State Transitions. Plant Physiology. 2020 Aug 1;183(4):1749–64. Available from: 10.1104/pp.20.00384

30. Nawrocki WJ, Liu X, Raber B, Hu C, de Vitry C, Bennett DIG, et al. Molecular origins of induction and loss of photoinhibition-related energy dissipation qI. Sci Adv. 2021 Dec 24;7(52):eabj0055.

31. Allorent G, Tokutsu R, Roach T, Peers G, Cardol P, Girard-Bascou J, et al. A Dual Strategy to Cope with High Light in Chlamydomonas reinhardtii. The Plant Cell. 2013 Feb;25(2):545–57.

32. Ruiz-Sola MÁ, Petroutsos D. A Toolkit for the Characterization of the Photoprotective Capacity of Green Algae. In: Maréchal E, editor. Plastids. New York, NY: Springer US; 2018. p. 315–23.

33. Tietz S, Hall CC, Cruz JA, Kramer DM. NPQ(T): A chlorophyll fluorescence parameter for rapid estimation and imaging of non-photochemical quenching of excitons in photosystem-II-associated antenna complexes. Plant, Cell & Environment. 2017;40(8):1243–55.

34. Nilkens M, Kress E, Lambrev P, Miloslavina Y, Müller M, Holzwarth AR, et al. Identification of a slowly inducible zeaxanthin-dependent component of non-photochemical quenching of chlorophyll fluorescence generated under steady-state conditions in Arabidopsis. Biochimica et Biophysica Acta (BBA) - Bioenergetics. 2010 Apr;1797(4):466–75.

35. Ramakers LAI, Harbinson J, Wientjes E, van Amerongen H. Unravelling the different components of nonphotochemical quenching using a novel analytical pipeline. New Phytologist [Internet]. [cited 2024 Dec 3];n/a(n/a). Available from: https://onlinelibrary.wiley.com/doi/abs/10.1111/nph.20271

36. Steen CJ, Burlacot A, Short AH, Niyogi KK, Fleming GR. Interplay between LHCSR proteins and state transitions governs the NPQ response in Chlamydomonas during light fluctuations. Plant, Cell & Environment. 2022;45(8):2428–45.

37. Harris PD, Eliezer NB, Keren N, Lerner E. Multiparameter-based photosynthetic state transitions of single phytoplankton cells bioRxiv; 2024. p. 2023.12.31.573751. Available from: https://www.biorxiv.org/content/10.1101/2023.12.31.573751v2

38. Mairal J, Bach F, Ponce J, Sapiro G. Online dictionary learning for sparse coding. In: Proceedings of the 26th Annual International Conference on Machine Learning. Montreal Quebec Canada: ACM; 2009. p. 689–96.

39. Fisher RA. The Use of Multiple Measurements in Taxonomic Problems. Annals of Eugenics. 1936;7(2):179–88.

40. Lahlou A, Tehrani HS, Coghill I, Shpinov Y, Mandal M, Plamont MA, et al. Fluorescence to measure light intensity. Nat Methods. 2023 Dec;20(12):1930–8. Available from: https://www.nature.com/articles/s41592-023-02063-y

41. Accueil ChlamyStation database. Available from: http://chlamystation.free.fr/

42. Depège N, Bellafiore S, Rochaix JD. Role of Chloroplast Protein Kinase Stt7 in LHCII Phosphorylation and State Transition in Chlamydomonas. Science. 2003 Mar;299(5612):1572–5.

43. Bujaldon S. Les antennes photosynthétiques chez Chlamydomonas reinhardtiiL: biogénèse, fonction et régulation. Sorbonne université; 2023. Available from: https://theses.fr/2023SORUS346

44. Gorman DS, Levine RP. Cytochrome f and plastocyanin: Their sequence in the photosynthetic electron transport chain of Chlamydomonas reinhardi. Proceedings of the National Academy of Sciences. 1965 Dec;54(6):1665–9.

45. Chouket R, Pellissier-Tanon A, Lahlou A, Zhang R, Kim D, Plamont MA, et al. Extra kinetic dimensions for label discrimination. Nature Communications. 2022 Mar;13(1):1482.

46. Katz DS. The Streak Plate Protocol. Microbe Library [Internet]. 2008 [cited 2023 Jul 14]; Available from: https://asm.org:443/Protocols/The-Streak-Plate-Protocol

47. Piccioni RG, Chua N, Bennoun P. A Nuclear Mutant of *Chlamydomonas reinhardtii* Defective in Photosynthetic Photophosphorylation: Characterization of the Algal Coupling Factor ATPase. European Journal of Biochemistry. 1981 Jun;117(1):93–102. Available from: https://febs.onlinelibrary.wiley.com/doi/10.1111/j.1432-1033.1981.tb06307.x

48. Bonente G, Ballottari M, Truong TB, Morosinotto T, Ahn TK, Fleming GR, et al. Analysis of LhcSR3, a Protein Essential for Feedback De-Excitation in the Green Alga Chlamydomonas reinhardtii. PLOS Biology. 2011 Jan;9(1):e1000577.

49. Atteia A, de Vitry C, Pierre Y, Popot JL. Identification of mitochondrial proteins in membrane preparations from Chlamydomonas reinhardtii. Journal of Biological Chemistry. 1992 Jan 5;267(1):226–34. Available from: https://www.sciencedirect.com/science/article/pii/S0021925818484839

50. Bentley JL. Multidimensional binary search trees used for associative searching. Commun ACM [Internet]. 1975 Sep 1 [cited 2023 Jul 18];18(9):509–17. Available from: https://dl.acm.org/doi/10.1145/361002.361007

51. Pedregosa F, Varoquaux G, Gramfort A, Michel V, Thirion B, Grisel O, et al. Scikit-learn: Machine Learning in Python. Journal of Machine Learning Research. 2011;12(85):2825–30.

52. Duda RO, Hart PE, Stork DG. Part 1: Pattern Classification and Scene Analysis. New York etc.L: J. Wiley&sons. Vol. 1. 1973.

53. Roach T, Na CS. LHCSR3 affects de-coupling and re-coupling of LHCII to PSII during state transitions in Chlamydomonas reinhardtii. Scientific Reports. 2017 Mar;7(1):43145.

54. Peers G, Truong TB, Ostendorf E, Busch A, Elrad D, Grossman AR, et al. An ancient light-harvesting protein is critical for the regulation of algal photosynthesis. Nature. 2009 Nov;462(7272):518–21.

55. Kress E, Jahns P. The Dynamics of Energy Dissipation and Xanthophyll Conversion in Arabidopsis Indicate an Indirect Photoprotective Role of Zeaxanthin in Slowly Inducible and Relaxing Components of Non- photochemical Quenching of Excitation Energy. Front Plant Sci. 2017 Dec 8;8. Available from: https://www.frontiersin.org/journals/plant-science/articles/10.3389/fpls.2017.02094/full

56. Malnoë A. Photoinhibition or photoprotection of photosynthesis? Update on the (newly termed) sustained quenching component qH. Environmental and Experimental Botany. 2018 Oct 1;154:123–33. Available from: https://www.sciencedirect.com/science/article/pii/S0098847218301862

57. Brooks MD. A Suppressor of Quenching Regulates Photosynthetic Light Harvesting. UC Berkeley; 2012. Available from: https://escholarship.org/uc/item/0th8p5zq

58. Tyystjärvi E, Aro EM. The Rate Constant of Photoinhibition, Measured in Lincomycin-Treated Leaves, is Directly Proportional to Light Intensity. Proceedings of the National Academy of Sciences of the United States of America. 1996 Apr;93:2213–8.

59. Tyystjärvi E. Chapter Seven - Photoinhibition of Photosystem II. In: Jeon KW, editor. International Review of Cell and Molecular Biology. Academic Press; 2013. p. 243–303. (International Review of Cell and Molecular Biology; vol. 300).

60. Girolomoni L, Cazzaniga S, Pinnola A, Perozeni F, Ballottari M, Bassi R. LHCSR3 is a nonphotochemical quencher of both photosystems in Chlamydomonas reinhardtii. Proceedings of the National Academy of Sciences. 2019 Mar;116(10):4212–7.

61. Finazzi G, Johnson GN, Dallosto L, Joliot P, Wollman FA, Bassi R. A zeaxanthin-independent nonphotochemical quenching mechanism localized in the photosystem II core complex. Proceedings of the National Academy of Sciences. 2004 Aug 17;101(33):12375–80. Available from: https://www.pnas.org/doi/abs/10.1073/pnas.0404798101

62. Rintamäki E, Martinsuo P, Pursiheimo S, Aro EM. Cooperative regulation of light-harvesting complex II phosphorylation via the plastoquinol and ferredoxin-thioredoxin system in chloroplasts. Proceedings of the National Academy of Sciences. 2000 Oct 10 ;97(21):11644–9. Available from: https://www.pnas.org/doi/abs/10.1073/pnas.180054297

63. Vink M, Zer H, Alumot N, Gaathon A, Niyogi K, Herrmann RG, et al. Light-Modulated Exposure of the Light- Harvesting Complex II (LHCII) to Protein Kinase(s) and State Transition in *Chlamydomonas reinhardtii* Xanthophyll Mutants. Biochemistry. 2004 Jun 1;43(24):7824–33. Available from: https://pubs.acs.org/doi/10.1021/bi030267l

64. Johnson X, Vandystadt G, Bujaldon S, Wollman FA, Dubois R, Roussel P, et al. A new setup for in vivo fluorescence imaging of photosynthetic activity. Photosynthesis Research. 2009 Oct;102(1):85–93.

65. Nawrocki WJ, Liu X, Croce R. Chlamydomonas reinhardtii Exhibits De Facto Constitutive NPQ Capacity in Physiologically Relevant Conditions. Plant Physiology. 2020 Jan;182(1):472–9.

66. Malerba ME, Palacios MM, Palacios Delgado YM, Beardall J, Marshall DJ. Cell size, photosynthesis and the package effect: An artificial selection approach. New Phytologist. 2018;219(1):449–61.

67. Cross FR, Umen JG. The chlamydomonas cell cycle. The Plant Journal. 2015;82(3):370–92.

68. McAdams HH, Arkin A. Stochastic mechanisms in gene expression. Proceedings of the National Academy of Sciences. 1997 Feb;94(3):814–9.

69. Elowitz MB, Levine AJ, Siggia ED, Swain PS. Stochastic Gene Expression in a Single Cell. Science. 2002 Aug;297(5584):1183–6.

70. Simpson EH. The Interpretation of Interaction in Contingency Tables. Journal of the Royal Statistical Society: Series B (Methodological). 1951;13(2):238–41.

71. Zhang R, Chouket R, Plamont MA, Kelemen Z, Espagne A, Tebo AG, et al. Macroscale fluorescence imaging against autofluorescence under ambient light. Light: Science & Applications. 2018 Nov;7(1):97.

72. Pech-Pacheco JL, Cristobal G, Chamorro-Martinez J, Fernandez-Valdivia J. Diatom autofocusing in brightfield microscopy: A comparative study. In: Proceedings 15th International Conference on Pattern Recognition ICPR- 2000. 2000. p. 314–7 vol.3.

73. Kosuge K, Tokutsu R, Kim E, Akimoto S, Yokono M, Ueno Y, et al. LHCSR1-dependent fluorescence quenching is mediated by excitation energy transfer from LHCII to photosystem I in Chlamydomonas reinhardtii. Proceedings of the National Academy of Sciences. 2018 Apr;115(14):3722–7.

74. Cantrell M, Ware MA, Peers G. Characterizing compensatory mechanisms in the absence of photoprotective qE in Chlamydomonas reinhardtii. Photosynth Res. 2023 Oct 1;158(1):23–39. Available from: 10.1007/s11120-023-01037-7

75. Alienor134. Alienor134/Image_segmentation: 2021.

76. Schreiber U. Pulse-Amplitude-Modulation (PAM) Fluorometry and Saturation Pulse Method: An Overview. In: Papageorgiou GC, Govindjee, editors. Chlorophyll a Fluorescence: A Signature of Photosynthesis. Dordrecht: Springer Netherlands; 2004. p. 279–319. (Advances in Photosynthesis and Respiration).

77. Virtanen P, Gommers R, Oliphant TE, Haberland M, Reddy T, Cournapeau D, et al. SciPy 1.0: fundamental algorithms for scientific computing in Python. Nat Methods 2020 Mar;17(3):261–72. Available from: https://www.nature.com/articles/s41592-019-0686-2

