## Supplementary_information for "Interplay between high-energy quenching and state transitions in *Chlamydomonas reinhardtii*: a single-cell approach"

July 16, 2025

**5 Comparison with *ad hoc* metrics** **18**

**6 Sources of heterogeneity** **19**

**7 Statistical tests** **20**
7.1 Monoclonal experiments . . . 20
7.2 Main text figures . . . 20

**8 Evaluation of the relevance of the axis $\tilde{q}\mathbf{I}$** **22**

### 1 Description of the fluorescence microscope

The home-made fluorescence microscope used for the experiments is pictured in Figure S1. In the following we present the technical choices for the illumination and the imaging parts, and the instrument controls. We built the microscope with Thorlabs parts, in three stages combined with thick Thorlabs posts to maximize the stability of the setup.

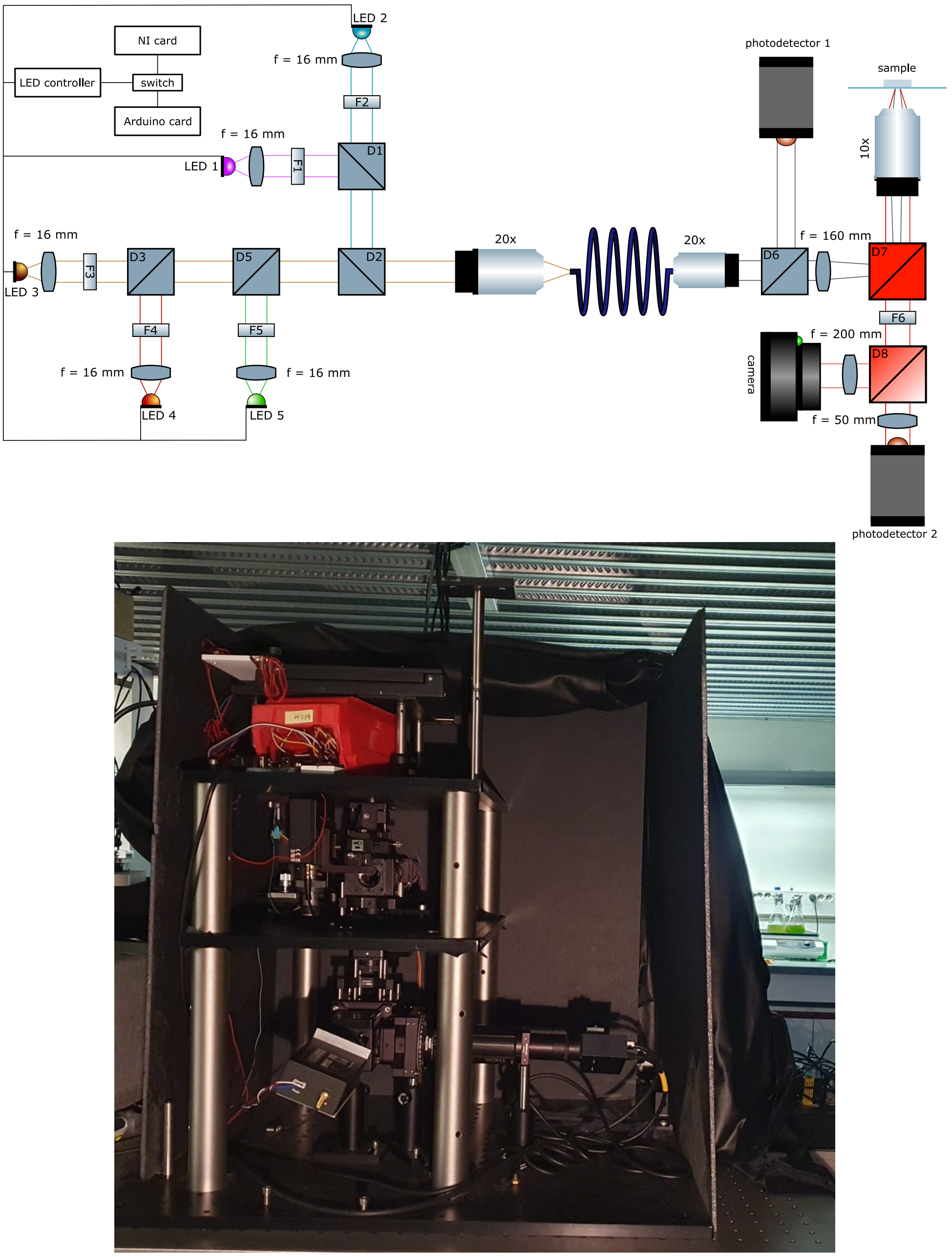

**Figure S1: Optical setup.** Top: Scheme of the optical setup. LEDs and band filters: 1: 405 nm, 2: 470 nm, 3: 690 nm, 4: 650 nm, 5: 550 nm; Dichroic high-pass filters: D1: 425 nm (to combine 405 nm and 470 nm), D2: 700 nm, D3: 665 nm (to combine 690 nm and 650 nm), D5: 560 nm; D6: glass plate to reflect incident light on detector, D7: 506 nm (to send light excitation towards the sample and collect the fluorescence), D8: 699 nm (cuts-off in the middle of the ChlF spectrum to split the signal between the photodetector and the camera - can be removed or replaced by a mirror); Bottom: Picture of the setup.

#### 1.1 Light part

The five light sources (Table S1) are high-power LEDs mounted on cages (*CXY1 - Thorlabs*). They are collimated by lenses with a focal distance of 16 mm (*ACL25416U - Thorlabs*) then filtered by band-pass filters (Table S1). A SWP 650 nm filter was added in front of the green LED L5 because the latter also emits in the red. The lights are combined by dichroic filters (Table S1) to be merged and injected into an optical fiber ($\emptyset = 400\ \mu m, NA = 0.5$ -

*M45L02 Thorlabs*) through a microscope objective (*Plan APO 20X/0.75 DIC N2 - Nikon*). A filter wheel (*FW102C - Thorlabs*) is set in front of the blue LED allowing to explore a range of intensity from 1 $\mu$mol(photons) $\cdot$ m$^{-2}$ $\cdot$ s$^{-1}$to 10000 $\mu$mol(photons) $\cdot$ m$^{-2}$ $\cdot$ s$^{-1}$. The LEDs are controlled by a voltage to intensity converter (*DC4100 - Thorlabs*). It can power four LEDs and is controlled by four voltage inputs. Depending on the type of signal expected, a different controller is used to send the voltage inputs, and a switch allows to keep the signals into two separate circuits. A National Instruments Data Acquisition Card *SCB68A* with *PCI 6374* generates time signals written as Python arrays, or an *Arduino Uno* controller generates precisely-timed pulses of light. The spectra of the blue and purple filtered LEDs used in this work have been acquired on a LPS 220 spectrofluorometer (PTI, Monmouth Junction, NJ) and correspond to 470±10 nm and 405±7 nm.

| **name** | **quality** | **wavelength** | **reference** | **supplier** |
|---|---|---|---|---|
| L1 | high power LED | purple | LHUV0-405-A065 | Lumileds |
| L2 | high power LED | blue | LXZ1-PB01 | Lumileds |
| L3 | high power LED | far-red | | Roithner Lasertechnik |
| L4 | high power LED | red | LXM3-PD01 | Lumileds |
| L5 | high power LED | green | | Lumileds |
| F1 | band-pass filter | 405/20 | ZET405/20x | Chroma |
| F2 | band-pass filter | 479/40 | FF01-479/40-25 | Semrock |
| F3 | band-pass filter | 690/8 | FF01 690/8 | Semrock |
| F4 | band-pass filter | 650/13 | FF01-650/13 | Semrock |
| F5 | band-pass filter | 550/15 | ET550/15x | Chroma |
| F6 | band-pass filter | 775/140 | FF01 775/140 | AHF |
| F6bis | band-pass filter | 675/90 | 675/90 ET | AHF |
| D1 | long-pass dichroic filter | 425 | t425lpxr | Chroma |
| D2 | long-pass dichroic filter | 700 | FF700-Di01 | Semrock |
| D3 | long-pass dichroic filter | 665 | FF665-Di02 | Semrock |
| D5 | long-pass dichroic filter | 560 | FF01 560 Di01 | Semrock |
| D6 | glass microscope plate | NA | | |
| D7 | long-pass dichroic filter | 506 | FF506-Di03 | Semrock |
| D8 | long-pass dichroic filter | 699 | FF699-FDi01 | Semrock |

**Table S1: Illumination equipment.** D7 can be replaced by D8 to excite the photosynthetic organisms with the green and red LEDs (fluorescence emission: 650-750 nm).

#### 1.2 Imaging part

The optical fiber output is connected to a microscope objective (*Plan 10x/.25 - Olympus*). A fraction of the illumination flux is reflected towards a photon counter by a microscope glass plate D6 installed on the light path. It allows to monitor the intensity of the LEDs for a given voltage input with Photodetector 1 (*Multi-Pixel Photon Counter C13366 - Hamamatsu*). The light path crosses a tunable diaphragm and a lens (*focal length 150 mm, AC254-150-A-ML - Thorlabs*). A dichroic filter D7 (Table S1) reflects the excitation light towards a microscope objective (*Objective Fluar 10x-0.50-M27 - Zeiss*). The objective can easily be changed for another magnification, with a flexibility in the working distance in the range of several centimeters, thanks to Thorlab rings stacking. The objective is mounted on a plate that moves in the $z$-direction to control the focus (*NFL5DP20/M - Thorlabs*). The plate can also be controlled by a piezoelectric (*KPZ101 - Thorlabs*, and we added the possibility to use a motor controlled by an Arduino to move the stage with a coarser precision[1]). The diaphragm is conjugated with the microscope plane and allows to reduce the diameter of the excitation light, and excite a limited area in the sample. The sample is mounted onto an automated platform with $x$ and $y$ movements[2]. The fluorescence is collected by the same microscope objective and is transmitted by the dichroic filter D8 (Table S1) and a fluorescence filter F6 (Table S1) towards the measuring devices. The fluorescence light can be either collected on a camera (*Ueye 3060CT-M - IDS*) conjugated with a tube lens (2 inches, focal length 200 mm, to maximally collect the photons), or a Photodetector 2 (*Multi-Pixel Photon Counter C13366 - Hamamatsu*) conjugated with a lens of focal length 50 mm. We can also collect the signal on both, depending on the filter used: mirror, dichroic filter D8 whose cut-off is in the middle

[1]https://github.com/SonyCSLParis/Motorized-stage
[2]https://github.com/SonyCSLParis/Motorized-stage/XY-stage

of the ChlF spectrum (Table S1), or no filter. The camera has adaptable gain and exposure time and is triggered internally or externally depending on the experiment. Unless specified, the camera was neither binned nor sub-sampled and the image size is 1936×1216 pixels$^2$.

#### 1.3 Heterogeneity of illumination

To study the homogeneity of the field of illumination, we performed the calibration process using Dronpa-2 solution as described in [1]. We analyzed the temporal response of the fluorescence of **Dronpa-2** extracted from the movie. Figure S2a shows a frame of the movie where the blue and purple LEDs are ON, while Figure S2b shows a slide where only the blue LED is on, evidencing the decay of fluorescence. The video ($608 \times 968$ pixel$^2$) was downsized by a $3 \times 3$ averaging kernel to improve signal-to-noise ratio. The 470 LED was kept constant and the 405 one was switched ON and OFF four times for 10 s every 24 s. The first response was discarded due to the difference of response of the dark state of Dronpa-2.[2] The mask of the illuminated area was evaluated with a threshold on the intensity image and the following operations were performed only on the unmasked pixels. Monoexponential fits were performed on each ON-OFF and OFF-ON phase and the retrieved characteristic times were averaged. From the extracted characteristic times, the light intensity in $\mu mol(photons).m^{-2}.s^{-1}$ was computed for each pixel using the molecular action cross-sections for photoisomerisation of Dronpa-2 [1]. The mean value of light intensity in the field in the given conditions is 8.0 and 1.4 $mmol(photons).m^{-2}.s^{-1}$ for the 470 and 405 LEDs respectively. The conditions chosen for these experiments are optimal for computing the distribution map with high resolution. In the experiments with algae, an optical density filter is placed in front of the 470 LED, before the injection in the optical fiber. The map of intensities derived from the relaxation times of the fluorescence evolution for the blue LED and the purple LED are shown in Figure S2c,d. The distribution of intensities presents a narrow distribution (Figures S2e,f). There is a 16% variation of light intensity in the field of view for the blue actinic LED and 30% variation for the purple SP LED.

#### 1.4 Instrument control

##### 1.4.1 Automated instrument control

The automated instruments (filter wheel, $x - z$ displacements, LED controller, LED switch) are controlled by serial communication with Python codes. The excitation signals are sent either through an Arduino Uno microcontroller or a DAQ National Instruments card (*SCB68A* with *PCI 6374*). The fluorescence signals are acquired through a camera (*UEye 3060CT-M - IDS*) controlled with Python or through the MPPC signals collected with the DAQ card.

##### 1.4.2 Autofocus routine

In the protocols described in subsection 3.1, experiments last several hours and the sample can move under the microscope and lose focus. With a motor driving vertical motion of the microscope stage and controlled by an Arduino, we operated an autofocus routine at low light before the dark adaptation, at each start of four consecutive reference protocol. The motor moves until the image captured presents a maximal Laplacian variance [3].

##### 1.4.3 Data management

For each experiment, the codes and the results are saved in a database MongoDB with the Python library Sacred.[3] The experiments are accessible with the interface Omniboard [4] which allows to compare the results.

[3]https://github.com/IDSIA/sacred
[4]https://github.com/vivekratnavel/omniboard

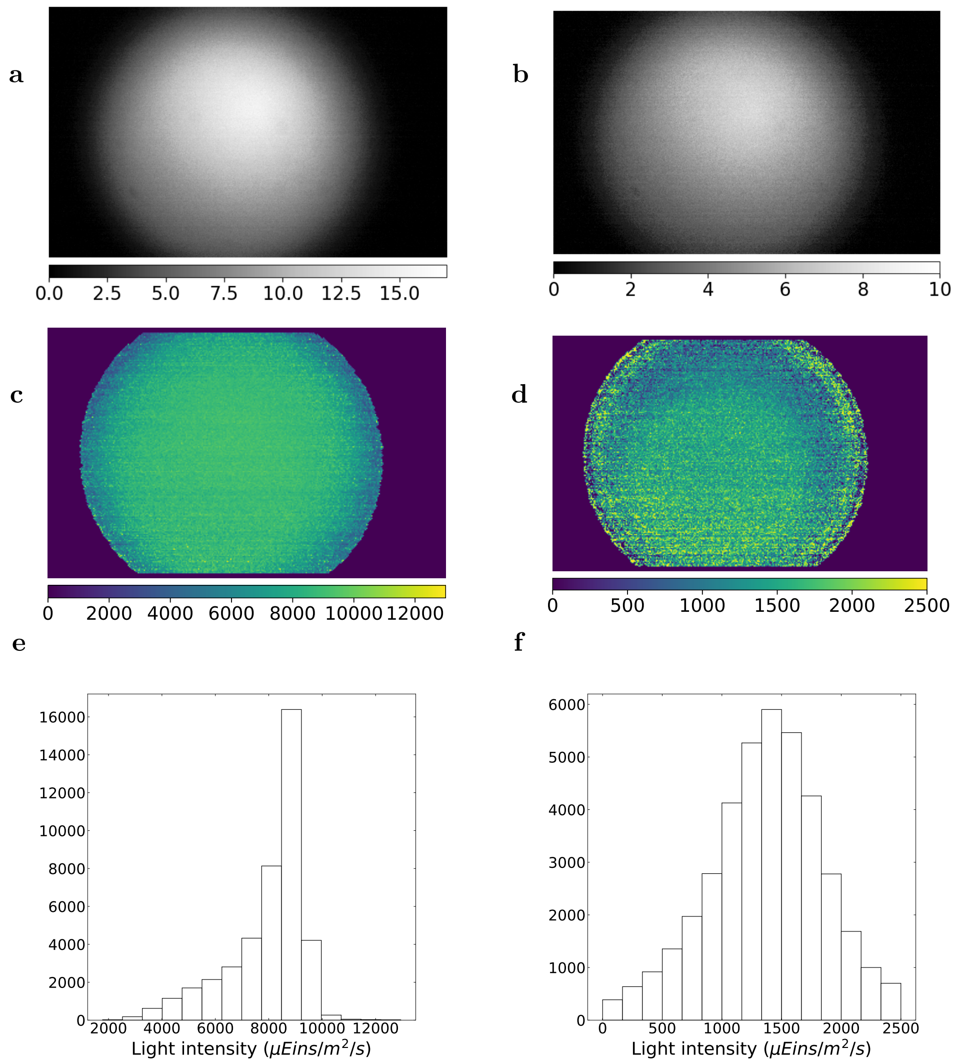

**Figure S2: Calibration of the light sources**. **a,b**: Intensity map of the 470 (resp. 405) LED in the focal plane calibrated with **Dronpa-2** for voltage inputs of the LED controller 500 mV (resp. 250 mV); **c,d**: Histogram of the intensity values, $(\mu, \sigma)$ = (8.0,1.3) (resp. (1.4,0.4) $mmol(photons).m^{-2}.s^{-1}$).

### 2 Biological samples

#### 2.1 Investigation of the strain *npq*4

The mutant *npq*4 lacks the LHCSR3 protein and it is commonly used in the literature as a negative control for qE. We noticed that the mutant *npq*4 exhibited qE after 4 h of HL-treatment under the microscope on the agarose pad. As shown in Figure S3b the fluorescence drops sharply at the light onset, characteristic of qE. The amplitude of the drop lies between the characteristic drops found in ***Pop_0*** (Figure S3a) and ***Pop_qE*** (Figure S3c). This behavior has previously been identified in the literature and attributed to an over-expression of LHCSR1 to compensate for the absence of LHCSR3[4, 5]. See however [6] for a different view on the function of LHCSR1.

Rather than using a double-mutant of LHCSR1 and LHCSR3, which exhibits compensatory physiological reorganization [7], we chose to grow cells in low light in the presence of acetate (see Methods) to create qE-free conditions. Conversely, for conditions requiring qE, cells were pretreated under HL after being transferred to acetate-free medium (see Methods).

#### 2.2 Evidence of a slow relaxing NPQ component

We formulated the hypothesis that the slow NPQ component corresponds to photoinhibition. To test this, we used 1 mM of lincomycin, an inhibitor of D1 protein repair. Indeed, PSII photodamage and qI occur when excessive light disrupts the balance of the continuous degradation and repair process of reaction centers involving the D1 protein, encoded in the chloroplast genome [8, 9]. Experiments were conducted using the SpeedZen set-up, as de-

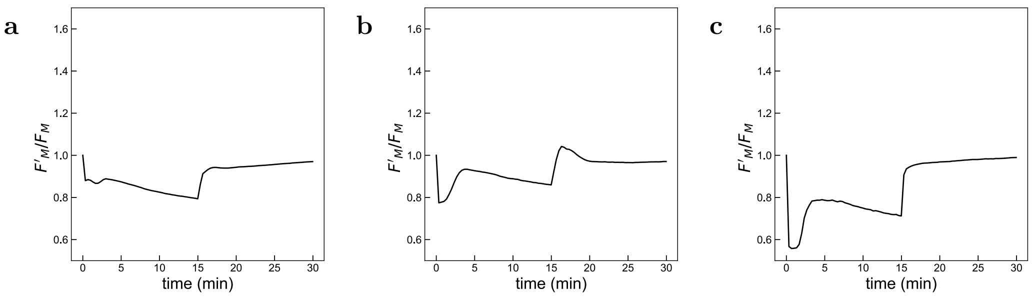

**Figure S3: Investigation of HL-treatment in npq4 strain.** Average $F'_m$ response of the population of algae in an agarose pad observed under the fluorescence microscope (normalized to $F_m$). **a**: ***Pop_0*** (*stt7-1* harvested from growth light); **b**: *npq4* after HL-treatment, fourth repeat of the reference protocol. **c**: ***Pop_qE*** (*stt7-1* after HL-treatement).

scribed in the Methods, to evaluate population's response to high-light illumination and its relaxation in the dark. We applied the basic excitation pattern on the SpeedZen, exposing a culture of stt7-1 to high-light (740 $\mu$mol(photons) $\cdot$ m$^{-2}$ $\cdot$ s$^{-1}$, 620 nm) for 15 min, followed by 15 min of dark relaxation, while measuring the evolution of $F'_m$ with saturating pulses (SP).

In our approach, we suggest that after two consecutive illuminations with the reference protocol, the D1 protein repair mechanism is active and the slow NPQ mechanism reaches steady-state. We tested this hypothesis by adding lincomycin in the dark adapted population just before the first illumination, and also after applying the light protocol two times in a row, and comparing the $F'_m$ traces to the control population without lincomycin.

In the first illumination (Figure S4a) we see no significant difference (P-value = 0.36) between the lincomycin-treated population and the control populations after 15 min of high light. However, at the end of the 15-minute dark relaxation we observe a significant (P-value = 0.001) but very small difference in the recovery. Suggesting that even without lincomycin, the D1 protein repair mechanism is only minimally active in the first illumination. After two consecutive illuminations (Figure S4b) we start noticing the presence of the fast NPQ component qE that is induced by the consecutive illuminations. But more importantly we see an increase in the slope of the slow NPQ component with the addition of lincomycin. Indeed, after 15 min of high light, $F'_m/F_m$ is significantly higher (P-value = 10-7) for the control without lincomycin (0.726 $\pm$ 0.003) compared to the lincomycin-treated population (0.68 $\pm$ 0.01). As expected, we notice a greater effect after 15 min of relaxation in darkness, with $F'_m/F_m$ values of 0.86 $\pm$ 0.01 for the control condition and 0.75 $\pm$ 0.01 for the lincomycin-treated sample (P-value = 10-8). These results support our hypothesis that by the third illumination, the D1 repair mechanism is fully active and compensates the high-light-induced PSII damage, leading to a reduced contribution of qI in the $F'_m$ traces.

To test whether the small effect of lincomycin in NPQ recovery after the first illumination protocol is indeed related to photoinhibition, we repeated the experiment with 40 min of relaxation in darkness instead of 15 min. Expecting this extended period would allow sufficient time for the D1 protein repair mechanism to activate and for the slow NPQ component to relax in the control sample. We also spaced each SP in darkness by 1 minute, rather than 20 s, to minimize any HL stress associated to SP. As can be seen in Figure S4c, after 40 min of dark relaxation we measured a significant difference (P-value = 10-7) in recovery between the lincomycin-treated population ($F'_m/F_m$ = 0.690 $\pm$ 0.004) and the control population without lincomycin ($F'_m/F_m$ = 0.756 $\pm$ 0.004). Therefore, we consider that the slowly reversible quenching in our dataset is dominated by qI, although a contribution from qZ cannot be strictly excluded [10].

#### 2.3 Growth of algae on the pad

We left the algae to grow with intermittent 15 min HL/15 min dark (HL: 400 $\mu$mol(photons) $\cdot$ m$^{-2}$ $\cdot$ s$^{-1}$, 470$\pm$10 nm) for several days and showed the growth was not limited by the sample preparation (Figure S5a). In Figure S5b we show the algae after a first division. Figure S5c shows the same sample after four and a half day. The surface occupied by algae was multiplied by 14 between the two images.

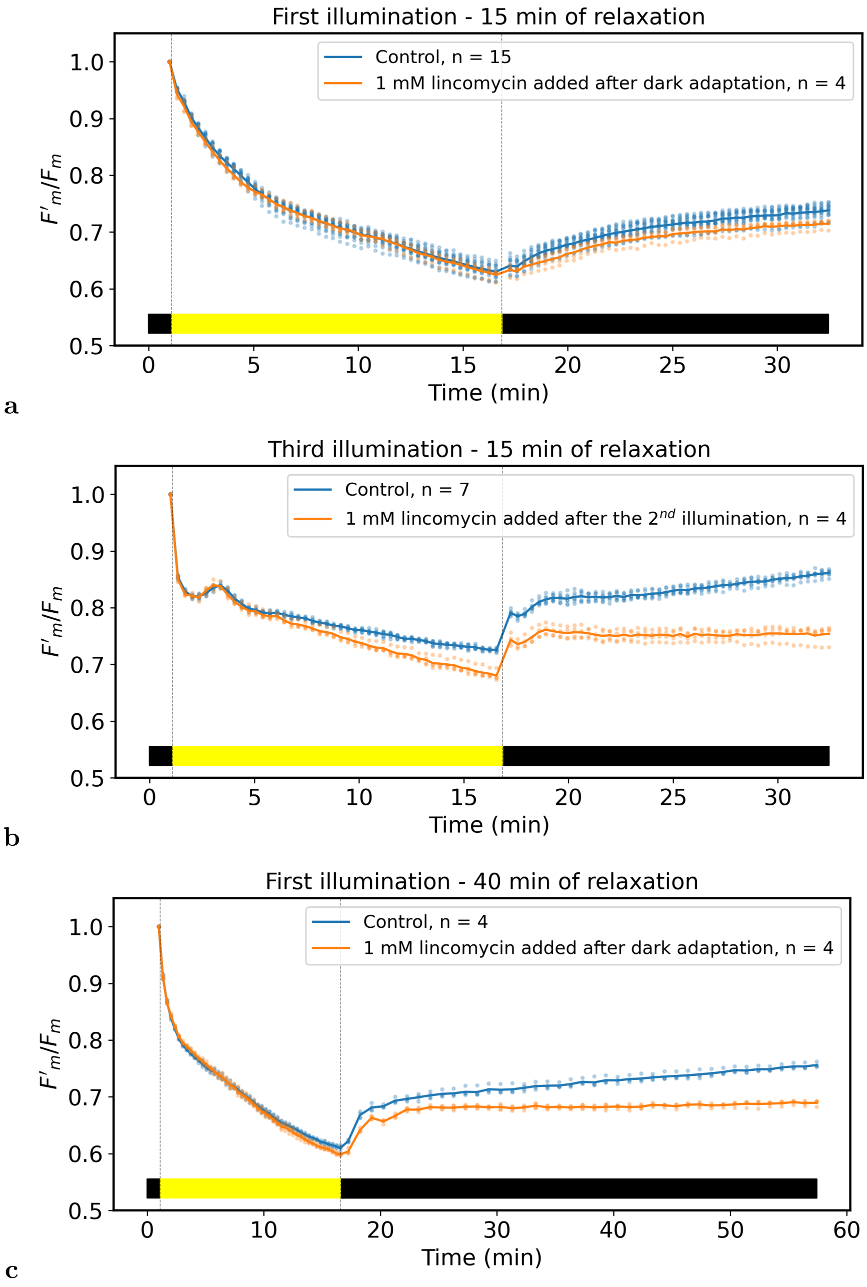

**Figure S4: Effect of lincomycin treatment on the slow NPQ component.** $F'_m$ response acquired on a population of stt7-1 on the SpeedZen exposed to lincomycin either after dark adaptation (a) and (c) or after the second illumination of the protocol (b). The dots represent experimental data from different technical replicates, while the continuous lines connect the averages across the replicates for each time point. High-light: 740 $\mu$mol(photons) $\cdot$ m$^{-2}$ $\cdot$ s$^{-1}$, 620 nm. Number of repeats: n (see legend).

#### 2.4 Image analysis of the single-cell Chlorophyll fluorescence movies

The algae in the video were segmented using local mathematical morphology operations described in *Chouket et al* [11] with the codes given in the corresponding Supplementary Materials [12] developed to segment fluorescent bacteria. As opposed to bacteria, algae are round and do not display a local maxima of fluorescence in the cell center. Instead, they show dark regions in the middle, corresponding to the part unoccupied by the fluorescent chloroplast. As a result, we changed the watershed source identification method: instead of locating local maxima of fluorescence, we locate the center of the algae. We begin by using a fill-holes operation to obtain only round and filled objects, which is not initially the case because of the dark areas. A simplified way to find the center of a disk with mathematical morphology operation is the following. Using a binary mask of the object, multiple consecutive erosions of one pixel are performed. After each erosion, a dilation is used to test if the center of the object has been reached. Since the disk is convex, an erosion followed by a dilation is fully reversible, except in one case. If the pixel is not reconstructed by the dilation, it means

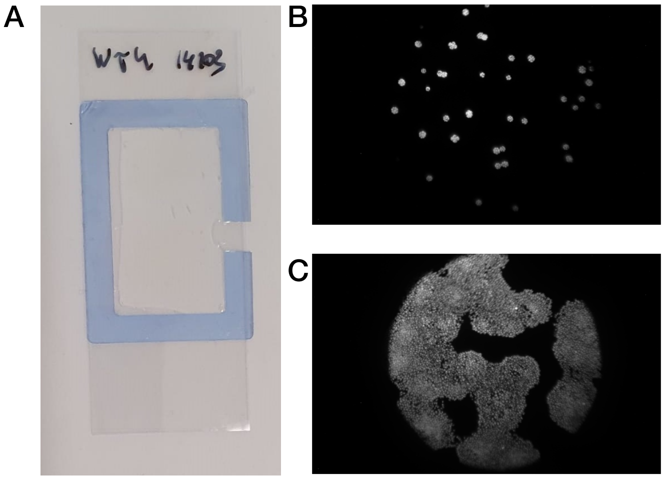

**Figure S5: Agarose pad. a**: Sample preparation: the medium mixed with agarose is spread on a pad and solidified. A drop of algae is deposited on the agar mix and covered by a microscope slide. **b,c**: Demonstration of the viability of the cells on the agarose pad. Fluorescence images of wild-type strain taken four and a half days apart. The surface occupied by the algae is multiplied by 14.

that the sphere has been completely eroded, and that the pixel was the furthest from the sphere boundaries, ie. the disk center. The number of steps it took to reach this non-return step is the distance of the center to the edge, here the radius of the algae in pixels. The results are displayed in Figure S6.

We manually annotated a single image in order to evaluate the performance of the segmentation. We notice that the algae out of focus are not identified by the segmentation, which is not a problem because in practice they would receive a different light intensity and emit a low fluorescence signal resulting in a poor signal-to-noise ratio. It resulted in an F1-score of 73%.[5]

For long-lasting experiments as presented in Figure S10, the autofocus allows to keep the algae in the focal plane, but they can undergo horizontal shifts. It results in different positions of the algae between two videos. The algae are restored to their original position during the data processing by recovering the displacement vector between the two images. We recover it using a registration method based on the images cross-correlation [11].

#### 2.5 Algae synchronization

To assess the quality of the synchronization described in Methods, we plotted the number of algae segmented in the field of view over the experiment time. Over a 20 h long experiment, we observed an initial plateau, a sharp increase in the cell number and another potential plateau (see Figure S7). When fitting with a sigmoidal curve the evolution of the number of algae, we found a characteristic time below 3h, demonstrating that a synchronous rhythm is imprinted in the algae division.

[5]To compute the F1-score for segmentation, we first calculate the true positives (TP, pixels segmented as algae in the ground truth and predicted mask), false positives (FP - pixels segmented as algae that belong to background), and false negatives (FN - pixels segmented as background that are algae in the ground truth). These values can be derived from the overlapping regions between the predicted and ground truth masks. The precision (P) is defined as the ratio of true positives to the sum of true positives and false positives: precision = TP / (TP + FP). The recall (R) is defined as the ratio of true positives to the sum of true positives and false negatives: recall = TP / (TP + FN). The F1 score is then calculated as the harmonic mean of precision and recall: $F1 - score = 2\frac{P \times R}{P+R}$.

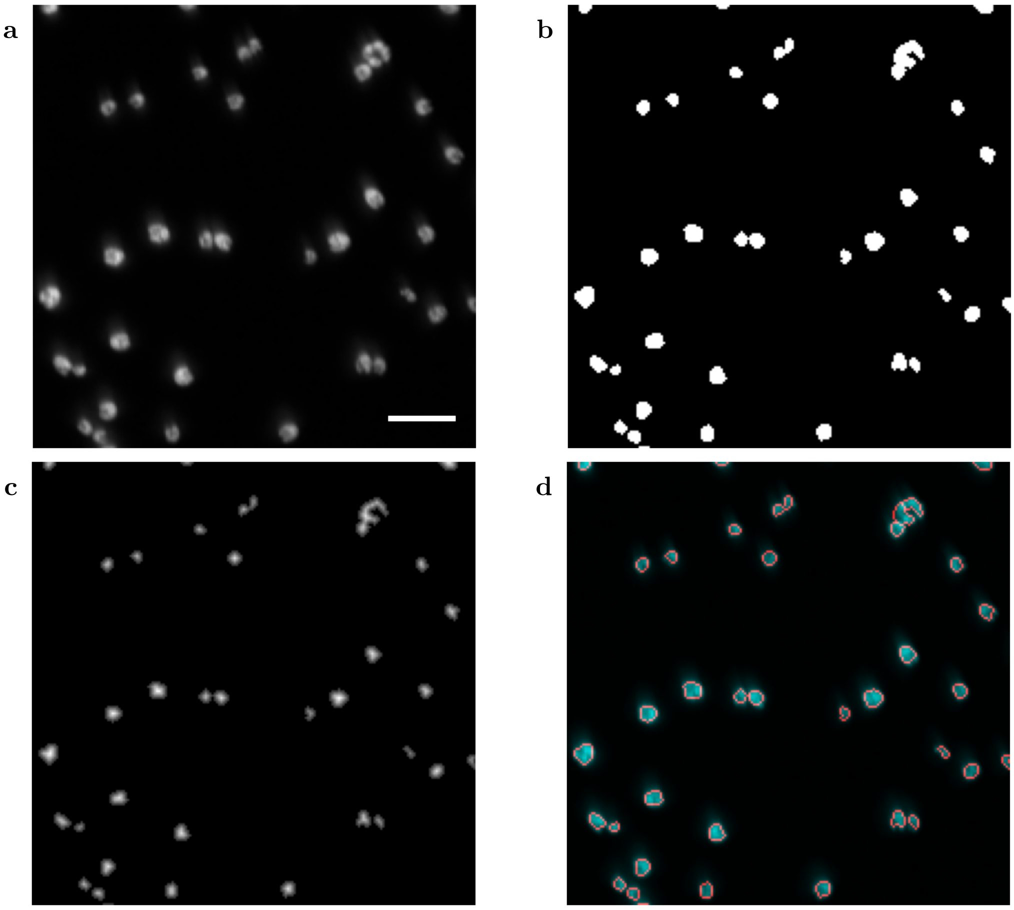

**Figure S6: Watershed segmentation of the fluorescence movies**. **a**: Grey-level image of the algae from a movie. Scale: 30 µm, cropped image; **b**: Intersection of the contrast mask and the auto-level mask, used as reference for the watershed segmentation; **c**: Distance map used to identify the local maxima; **d**: Contours of the segmented objects.

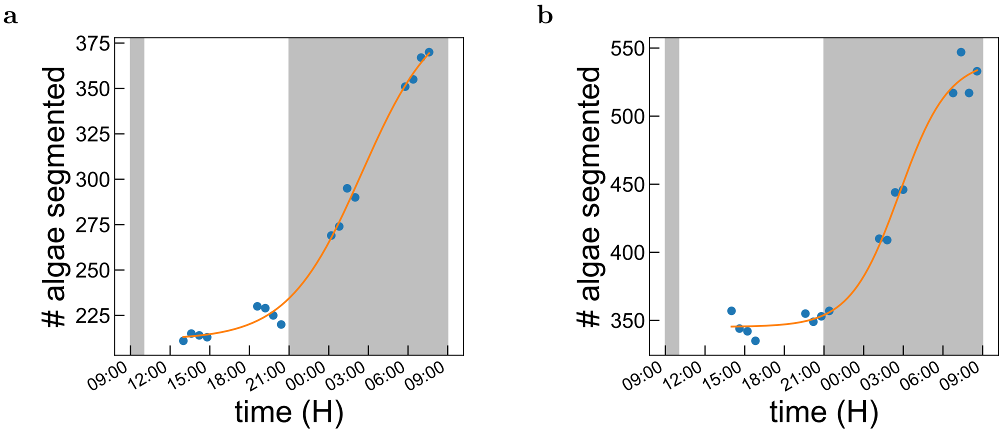

**Figure S7: Evaluation of the synchronicity**. Here the algae populations are the WT samples HL-treated under the microscope for the experiment described in Figure 4 of the Main Text. The algae are HL-treated under the microscope and taken out of the synchronization conditions, but the light-dark cycle to which they have been adapted is represented as dark areas in the figure. After a plateau during the expected light period, the algae start dividing at the onset of the expected dark period. The experiment is too short to assess that a plateau is reached after the division, but since the algae have been one day through HL-treatments and not in synchronized conditions, we can expect the synchronization to be lost after one day. **a,b**: number of algae segmented in the ChlF movie in the population $wt4a^{-}$ (resp. *cc*124) exposed to the light protocol under the microscope (see Main Text for sample conditions). A sigmoidal fit performed on the data points gives a characteristic time of 2h50 (resp. 1h50).

### 3 Data collection

#### 3.1 Illumination protocols

##### 3.1.1 Protocol inducing high-light stress and used to collect $F_m^{'}$ (reference protocol)

The reference protocol allows to induce simultaneously qE, qI and qT by applying a high light (400 $\mu$mol(photons) $\cdot$ m$^{-2}$ $\cdot$ s$^{-1}$, 470±10 nm) for 15 min and turning it off. This light is 10 times above the growth light (50 $\mu$mol(photons) $\cdot$ m$^{-2}$ $\cdot$ s$^{-1}$, white LED) corresponding to levels commonly used in the literature to induce light-stress response [13].

Simultaneously to the stressing light, we apply a saturating pulse every 20 s (1400 $\mu$mol(photons) $\cdot$ m$^{-2}$ $\cdot$ s$^{-1}$, 405±7 nm, 200 ms). The blue light is turned off when the saturating pulse is applied, this allows to remove the fluorescence contribution in response to the blue light. It allows to compare the fluorescence response to a saturating pulse during the light stress, but also in the dark when the blue light is turned off. Figure S8 shows the time evolution of the light level in the field of view.

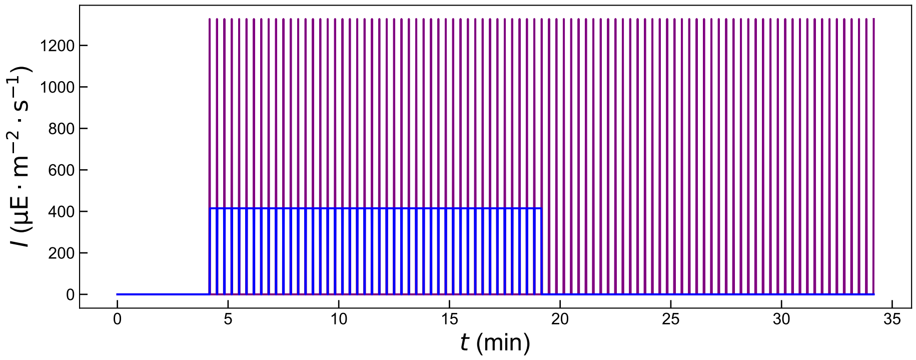

**Figure S8: Description of the reference protocol illumination.** The blue light induces the high-light stress (400 $\mu$mol(photons) $\cdot$ m$^{-2}$ $\cdot$ s$^{-1}$, 470±10 nm), the purple light pulses (1400 $\mu$mol(photons) $\cdot$ m$^{-2}$ $\cdot$ s$^{-1}$, 405±7 nm, 200 ms) saturate the photosynthetic chain to probe the fluorescence level under high-light and in the dark.

##### 3.1.2 Summary of the light protocols

To complement the description in the methods section, the light protocols are summarized in Table S2. The typical excitation sequence of the protocol allowing to generate the training set is displayed in Figure S10a.

**Table S2: Summary of the light protocols.** The actinic light is a LED (470±10 nm), the saturating pulses are produced with a LED (1400 $\mu$mol(photons) $\cdot$ m$^{-2}$ $\cdot$ s$^{-1}$, 405±7 nm, 200 ms) applied every 20 s.

| Name | Duration | Actinic light levels ($\mu$mol(photons) $\cdot$ m$^{-2}$ $\cdot$ s$^{-1}$) | **SP** | Figure |
|---|---|---|---|---|
| dark adaptation | 15 min | 0 | OFF | S10 |
| reference protocol | 30 min | 400 (470 nm) | ON | S8 |
| four reference protocol repeats | 2h | 400 (470 nm) | ON | S10 |
| HL-treatment | 1h20-4h | 400 (470 nm) | OFF | S10 |
| low-light relaxation | 45 | 40 (470 nm) | OFF | S10 |

#### 3.2 Evaluation of the quenching effect of the saturating pulses

In order to probe the fluorescence NPQ response under high-light and in the dark, we used a high saturating pulse frequency (1400 $\mu$mol(photons) $\cdot$ m$^{-2}$ $\cdot$ s$^{-1}$, 405±7 nm, 200 ms) to have a better temporal resolution of the stress-responses, at the risk of provoking a stress related to the saturating pulse. Although actinic light already saturates and provokes stress, we assessed if the saturating pulses significantly affected the fluorescence response by applying the reference protocol with actinic blue light set to zero (Figure S9b) as opposed to the reference protocol illustrated in Figure S9a. Cells of *stt7-1* taken from growth light and transferred to MIN medium as indicated in Methods were exposed to this SP assessment sequence after the 15 min dark adaptation followed by the four reference protocol repeats. The time evolution of the fluorescence NPQ response to the SP assessment pattern is plotted in Figure S9d, whereas the response to the last repeat of the reference protocol that was applied previously is plotted in Figure S9c. The condition of this population corresponds to ***Pop_0*** and is expected to present minimal NPQ response. The stress-response in Figure S9c is still much reacher with blue light on, demonstrating that the kinetics of ChlF response we observe mainly come from the light-stress provoked by the blue light, and not from the light-stress provoked by the saturating pulses. The slight increase of the response in Figure S9d may reflect the relaxation of the slowly-induced light stress response that we attribute to qI.

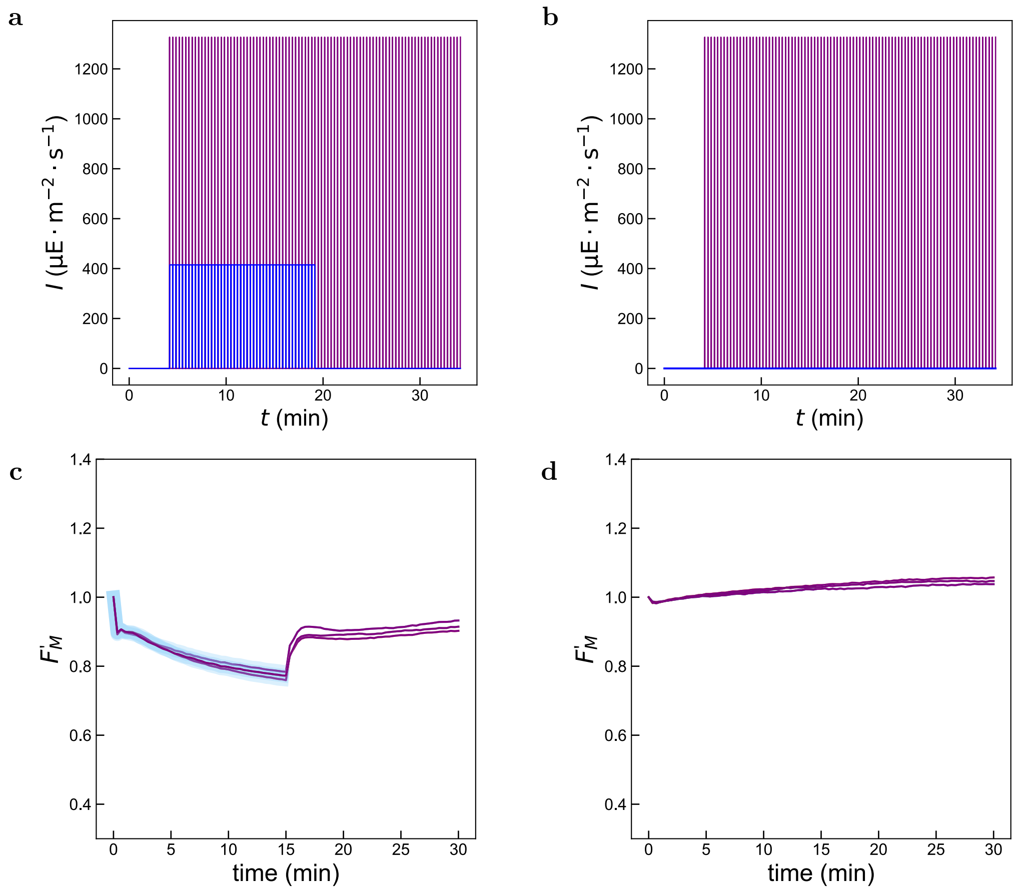

**Figure S9: Evaluation of the quenching effect of the saturating pulse**. **a**: Excitation sequence to investigate the light-stress induced by the saturating pulse when the blue light is turned on (reference protocol). **b**: Excitation sequence to investigate the light-stress induced by the saturating pulse when the blue light is turned off (SP assessment). **c**: $F'_m$ response of a *stt7-1* population to the last repeat of the four reference protocol repeats, before HL-treatment. Equivalent to ***Pop_0***. **d**: $F'_m$ response of a *stt7-1* population to a SP assessment after 15 min dark adaptation and four reference protocol repeats.

#### 3.3 Generation of the $F'_m$ traces

The reference dataset was generated by applying the full protocol described in S10a to the populations *stt7-1* and *wt4a*$^{-}$. The summary of these conditions is provided in Table 1 of the Main text. Figure S10a shows the evolution of the blue light intensity received by the algae through the different steps, without the saturating pulses. There are no saturating pulses during the HL-treatment and the relaxation, only during the reference protocol. Figure S10b,c show the fluorescence response to the consecutive light protocols for the two strains investigated (*wt4a*$^-$ and *stt7-1*). The reference protocol fluorescence response used to build the reference dataset as described in the Main Text are highlighted in the corresponding colors.

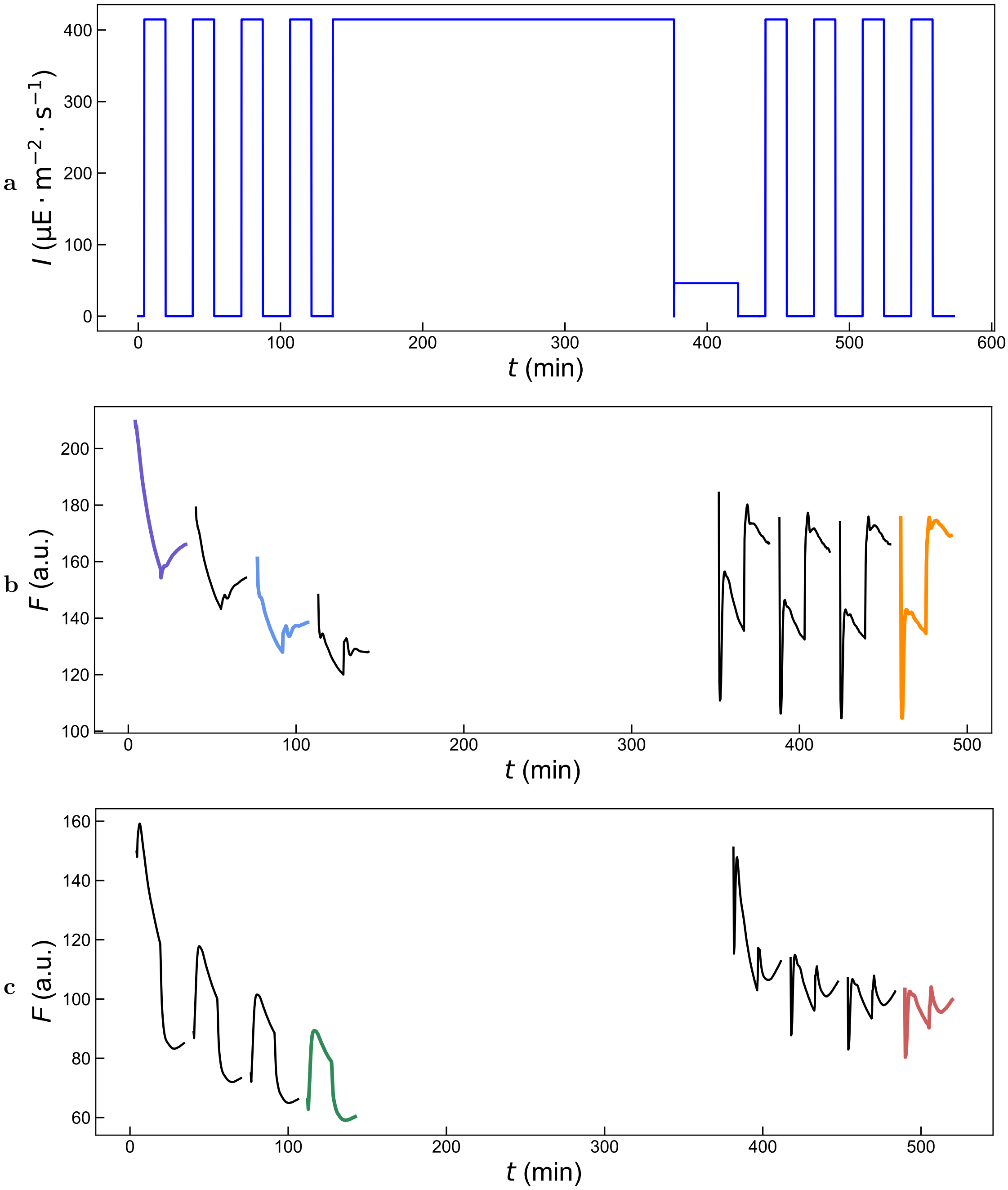

**Figure S10: Full excitation sequence allowing to probe qE, qI, qT, activate qE and achieve a negligible contribution of qI. a**: Blue light inducing the high-light stress (saturating purple light pulses not displayed), the HL-treatment and the photoinhibition; **b,c**: Response of *stt7-1* (resp. *wt4a*$^-$) to the full protocol, average over the fluorescence of the algae in the videos. The timescale does not correspond exactly to the theoretical pattern because the video processing is integrated in the experiment and one experiment needs to terminate before another starts. The colors correspond the the reference protocols response selected to build the dataset. Purple: ***Pop_qI***, blue: ***Pop_0***, orange: ***Pop_qE***, green: ***Pop_qT***, red: *wt4a*$^-$.

The reference protocol was repeated four times in a row before and after HL-treatment to reduce the contribution of qI. We see that after the third repeat, the final $F_m'$ is close to $F_m$ for *stt7-1* after HL-treatment, and for *wt4a*$^-$ before and after HL-treatment. We selected these three types of responses to collect the $F_m'$ traces of the populations ***Pop_qE***, ***Pop_qT*** and *wt4a*$^-$ respectively. For *stt7-1* before HL-treatment, we used the response to the first pattern to collect the $F_m'$ traces of the population ***Pop_qI***. We notice that the exposure to the reference protocol induces rapidly qE, as seen by the decrease of fluorescence response between two first pulses at the onset of the reference protocol. In order to minimize the effect of qE in this population of the dataset which is expected to show no stress response (***Pop_0***), we selected the third and not the fourth repeat of the reference protocol.

We also acquired similar data with monoclonal populations and *cc*124 populations that were not included in the dataset but used for investigation of the origin of the distribution of the NPQ scores (see Main Text).

For data processing, the $F'_m$ traces, response to saturating pulses only, were isolated from the whole fluorescence trace by selecting one in every 20 frames (framerate: 1 Hz, saturating pulse applied every 20 s). The data normalization is discussed in Section 4.

The dataset as well as a description of how to use it are available online [6]

[6] https://github.com/DreamRepo/NPQScore-data

### 4 Dictionary learning to represent the $F'_m$ traces

#### 4.1 Description of the Dictionary Learning hyperparameters

As presented in the Methods section, the dictionary learning aims at reconstructing a signal from a learned set of atoms. This set is learned by minimizing the loss function given in Eq.(1). The optimized metric is the reconstruction fidelity with a penalty on the sparsity level.

$$\min_{\mathbf{D},\mathbf{X}} \frac{1}{2}||\mathbf{Y} - \mathbf{D}\mathbf{X}||_2^2 + \lambda||\mathbf{X}||_1, \quad (1)$$

Where:

- $\mathbf{Y}$ is the input data matrix of dimension 91× $n_{samples}$,
- $\mathbf{D}$ is the dictionary matrix with $N_D$ atoms of dimension 91(normal matrix),
- $\mathbf{X}$ is the coefficient matrix of dimension $N_D$,
- $||\mathbf{Y} - \mathbf{D}\mathbf{X}||_2^2$ is the squared Euclidean norm of the reconstruction error,
- $||\mathbf{X}||_1$ is the L1 norm (sum of absolute values) of the coefficients,
- $\lambda$ is the regularization parameter that controls the sparsity level.

#### 4.2 Investigation of the hyperparameter space

We explored several parameters, based on the analysis of the reconstruction error.

- **Normalization method**: We tried normalizing $F'_m$ by
  - the standard deviation $\sigma = \sqrt{\frac{1}{N-1}\sum_{i=1}^{N}(F'_m(i) - \bar{F'_m})^2}$, a classical reduction method;
  - the first pulse value $F'_m(0)$ (also referred to as $F_m$);
  - the sum $\mathrm{S} = \sum_{i=1}^{N} F'_m(i)$ to obtain a probability law.
- **Dictionary Size** $N_D$: The size of the dictionary determines the number of atoms to be learned. The higher the size, the better the reconstruction. We explored sizes of dictionaries from 3 to 30 atoms.
- **Sparsity Level** $\lambda$: The sparsity enforces to learn to reconstruct the traces by combining a minimal number of atoms of the Dictionary. We explored sparsity constraints between $10^{-10}$ and $10^{-2}$.
- **Training set size** $n_{samples}$: This parameter corresponds to the number of $F'_m$ traces representing each class in the training. We used a balanced dataset, by selecting $n_{samples}$ in each class. The total training size is $4 \times n_{samples}$ because we have four classes. We tested values of $n_{samples}$ between 3 to 300 (representative of the size of the smallest class in the reference dataset).

Figure S11a and b show a poor and good reconstruction of a traces from HL-treated $wt4a^{-}$ population with a dictionary of 2 and 10 atoms respectively. Several number of training-set sizes, number of atoms and sparsity levels were explored to see their influence on the reconstruction error. We began by a grid search, splitting the exploration space by choosing seven values each for $N_D$, $\lambda$ and $n_{samples}$. Then, we refined the size of the search space around the optimum.

We selected the normalization by S after observing that it systematically yielded the lowest reconstruction error compared to the other normalization methods (Figure S11c). In the same Figure, we identify that the reconstruction error decreases with the number of atoms in the dictionary. In practice, when using the normalization by the sum $S$, and a dictionary larger than 12 atoms, some atoms were never used in the reconstruction of the training set, and were therefore irrelevant to train the LDA. This was true for all the values of $\lambda$ we investigated ($10^{-10}$ to $10^{-2}$). It is why the curve of the normalization by S stops at 12. The Figures S11d and S11e correspond to the results of the investigation limited to the normalization by S for clarity.

We investigated the impact of the size of the training set on the reconstruction error. Figure S11d shows that it is much less significant than the number of atoms. It shows that even a low amount of training data is sufficient to encompass the dynamics of the NPQ components in the dictionary. Therefore, the method could be generalized to studies at the population level with only a few repeats. Based on the curves, we selected 10 atoms for our dictionary, with a corresponding reconstruction error threshold of $2{\cdot}10^{-4}$.

Finally, we investigated the role of the parameter $\lambda$ on the reconstruction, as shown in Figure S11e (normalization by S, number of atoms: 10). Given the span of the error axis, we can deduce that it is minimal. It does however play a role on the number of atoms used for the reconstruction as we observed experimentally. We selected $\lambda = 10^{-6}$ to give maximal weight to the sparsity constraint in the metric optimization.

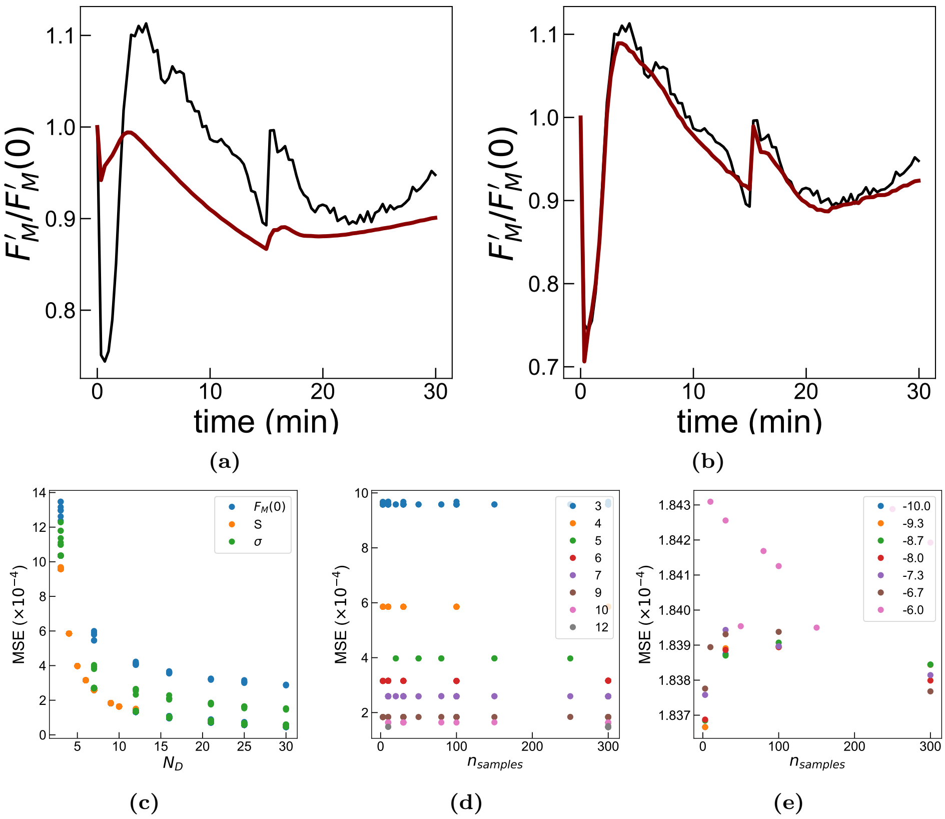

**Figure S11: Exploration of the role of the hyperparameters of the Dictionary Learning on the reconstruction error.** **a**: Fluorescence trace (black) and trace reconstructed with the dictionary learning method (red), with $N_D = 2$, using the normalization by $F'_m(0)$ and $n_{samples} = 300$. The mean squared error equals $6.7{\times}10^{-3}$ and $r^2 = -0.13$. **b**: Fluorescence trace (black) and trace reconstructed with the dictionary learning method (red), with $N_D = 10$, using the normalization by $S$ and $n_{samples} = 300$. The mean squared error equals $2.6{\times}10^{-4}$ and $r^2 = 0.95$. **c**: Impact of the normalization method and the number of atoms $N_D$ on the reconstruction error. Legend: normalization method; **d**: Impact of the training set size $n_{samples}$ and the number of atoms $N_D$ for dictionaries built using the normalization by $S$. Legend: number of samples per population (balanced dataset). **e**: Impact of the training set size $n_{samples}$ and the sparsity level $\lambda$ for dictionaries built using the normalization by $S$ and 10 atoms. Legend: $\log(\lambda)$.

#### 4.3 Validation of the reconstruction on $F'_m$ traces

We evaluated the performances of the first dimension reduction step (dictionary learning method) at reconstructing new data. First we investigated its capacity at reconstructing a validation set: monoclonal populations equivalent to ***Pop__0***, ***Pop__qE***, ***Pop__qT*** and ***Pop__qI*** not present in the training set. In figure S12b-e, we show traces overlayed with the dictionary reconstruction (seven traces with the poorest reconstruction from each class). We evaluate the quality of the reconstruction by computing the mean-squared error (MSE) and plot the MSE for each class in Figure S12j. We can identify a maximal average reconstruction error for the class ***Pop__qE*** at MSE $= 1.10^4$. In Figure S12f-i, we plot the reconstruction error for experimental datasets that were absent from the training set, at the fourth repeat, before and after HL-treatment. They consist of two wild-type populations, wild type $wt4a^-$ and wild type *cc124*. All these strains are expected to display qT, and qE when they are HL-treated. In all cases, the MSE is contained within the range of error of the training set (Figure S12k), therefore we can conclude that the dictionary reconstruction generalizes well to new data that combine the training traits. It serves as validation of the initial hypothesis that a fluorescence trace of combined traits is well represented as a linear combination of the fluorescence traces corresponding to individual traits.

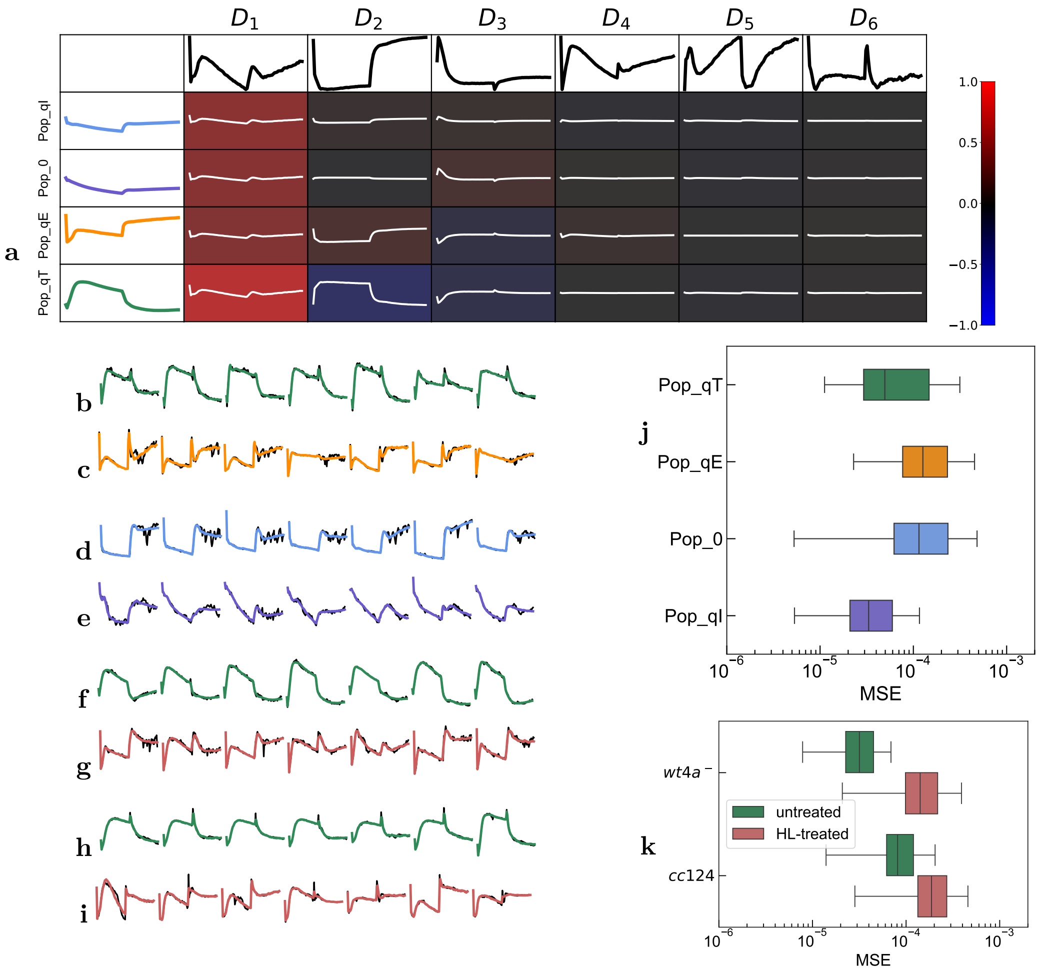

**Figure S12: Dictionary learning reconstruction for phenotypes absent from the training set.** **a**: The atoms $D_i$ are the learned representation of the dataset. Each element of the dataset can be reconstructed as a linear combination of the atoms $D_i$. In the table, the first column shows the reconstruction of the average trace of each population in the reference dataset. In each table cell, the color scales $Z_i$ while the trace is $Z_i \times D_i$; **b-e**: Raw (black) and reconstructed (colors) traces of the monoclonal populations not seen by the training set with the dictionary. The seven reconstructions with the highest Mean-Squared Error (MSE) are displayed. **b**: ***Pop_qT***, **c**: ***Pop_qE***, **d**: ***Pop_0***, **e**: ***Pop_qI***; **f-i**: Raw (black) and reconstructed ChlF traces of populations absent from the training set (except ***Pop_qT*** in **f**). The ChlF traces are measured before (green) and after (red) HL-treatment for 2-4 h. **f,g**: *wt4a*$^-$, **h,i**: **cc124** ; **j**: Distribution of the reconstruction mean squared error from monoclonal experiments not seen by the training set. ***Pop_qT***: 180 algae, ***Pop_qE***: 380 algae, ***Pop_0***: 317 algae, ***Pop_qI***: 326 algae; **k**: Distribution of the reconstruction mean squared error from experiments displayed in **f-i** *wt4a*$^-$: 196 algae, *cc*124: 164 algae. The traces plotted in **b-i** have the highest reconstruction error among all the algae in the experiment.

#### 5 Comparison with *ad hoc* metrics

To validate the method, we used *ad hoc* features identified in the literature allowing to derive quantitative values to represent the stresses. We compare the so-called relative NPQ values to the axis values and evaluated their correlation.

To evaluate the NPQ of a $F'_m$ trace of saturating pulses $F'_m$, the formula described in the literature is $NPQ(t_n) = \frac{F_m(0)-F_m(t_n)}{F_m(t_n)}$. We compare the NPQ($t_n$) values against the metrics we extracted with the machine learning pipeline in order to see if our findings are consistent with the literature. The fluorescence value at $t_n$ is highlighted on the top of the graphs, over the average traces of the training set overlaid. Several characteristic phases of the $F'_m$ traces have been identified to reflect either qE, qT and qI, based on mutants experiments [14–16]. We attributed time values of interest $t_n$ based on the local extrema described in [14] to identify them in each experiment. As described in the main text:

- The sharp decay of fluorescence at the light onset has been attributed to qE [14] and corresponds to NPQ($t_1$) (first SP after 10 s of HL). We compare it to the values computed for q̃E for all the algae in the class ***Pop__qE*** of the dataset and display the results in figure S13a.
- The gradual increase of the fluorescence under the light has been attributed to the migration of light-harvesting antennas to PSII and corresponds to NPQ($t_{45}$) (last SP after 15 min of HL). We compare it to the values computed for q̃T for all the algae in the class ***Pop__qT*** of the dataset and display the results in Figure S13b.
- The effect of qI contributes to a non-reversible decay of the fluorescence throughout the whole experiment (NPQ($t_{91}$) - last SP after 15 min in the dark following the 15 min of HL). We compare it to the values computed for q̃I for all the algae in the class ***Pop__qI*** of the dataset and display the results in Figure S13c.

In all cases we find a high correlation between the two scores for the class of interest.

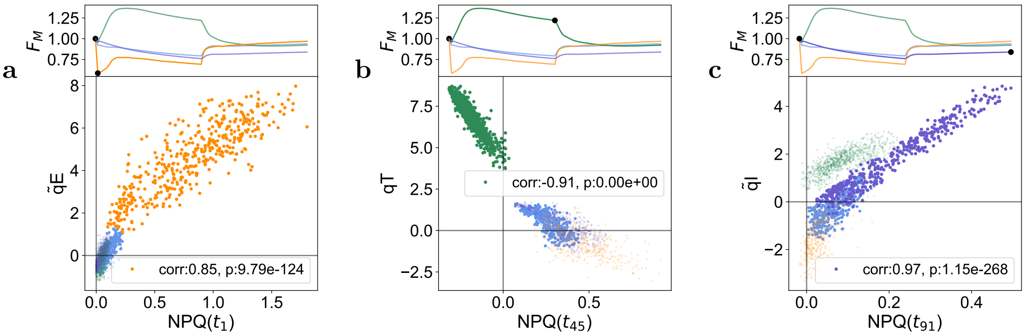

**Figure S13: Comparison of the NPQ features identified in the literature with the metrics extracted with the machine learning pipeline**. The average fluorescence traces of the training set are displayed on top, while the NPQ metrics are plotted against the scores computed with the machine learning pipeline. **a** shows the comparison of the effect of qE, contributing to the non-reversible decay of the fluorescence throughout the whole experiment (NPQ($t_1$)), with the values computed for q̃E for all the algae in the training set. The correlation coefficient displayed corresponds to the class ***Pop__qE*** (orange). **b** shows the comparison of the effect of qT, contributing to the non-reversible decay of the fluorescence throughout the whole experiment (NPQ($t_{45}$)), with the values computed for q̃T for all the algae in the training set. The correlation coefficient displayed corresponds to the class ***Pop__qT*** (green). **c** shows the comparison of the effect of qI, contributing to the non-reversible decay of the fluorescence throughout the whole experiment (NPQ($t_{91}$)), with the values computed for q̃I for all the algae in the training set. The correlation coefficient displayed corresponds to the class ***Pop__qI*** (purple).

### 6 Sources of heterogeneity

In order to test the correlation between the heterogeneity of illumination of the algae and the level of expression of qE, qT, and qI , we first exploited the calibration of the light intensity in the field of view with **Dronpa-2** (Figure S2). We then measured the NPQ-score of the monoclonal culture wild type $wt4a^-$ HL-activated over four reference protocol repeats. As displayed in Figure S14a–c, no significant correlation appears.

In order to test the contribution of the size of the algae to the variance of the NPQ scores, we also extracted the projected surface of each alga exposed to the light as derived from the number of pixels corresponding to the segmented alga. As shown in Figure S14d–e, we did not evidence any significant correlation.

Eventually, we evaluated if the spatial position of the algae could affect their NPQ score. Indeed, they may have been differently exposed to gas exchange from the sides of the observation chamber. As displayed in Figure S14g–i, we did not evidence any specific spatial pattern of the NPQ score.

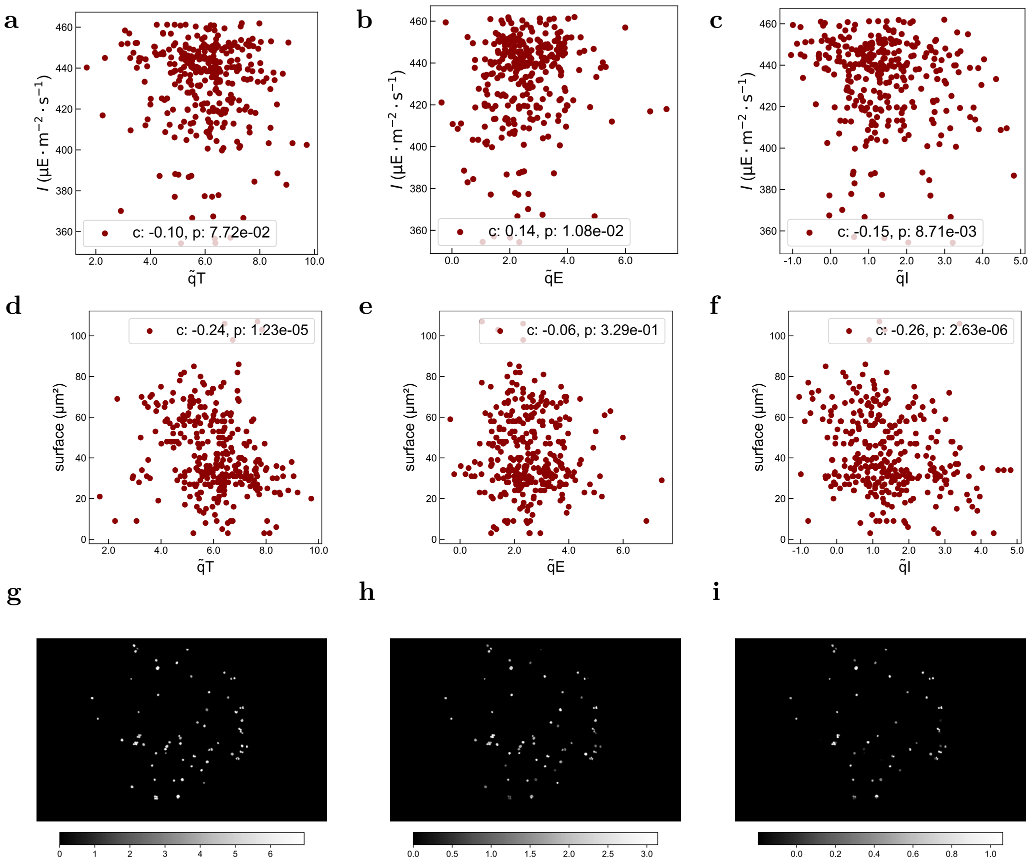

**Figure S14: Investigation of the correlation of the NPQ score with various parameters.** **a**–**c**: Dependence of the NPQ score ( **a**: q̃T; **b**: q̃E; **c**: q̃I) on light intensity; **d**–**f**: Dependence of the NPQ score ( **d**: q̃T; **e**: q̃E; **f**: q̃I) on the cell size; **g**–**i**: Dependence of the NPQ score ( **g**: q̃T; **h**: q̃E; **i**: q̃I) on the cell position in the observation chamber. Strain: monoclonal $wt4a^-$ HL-activated, **a-f** show overlap of the four consecutive reference protocol responses.

### 7 Statistical tests

#### 7.1 Monoclonal experiments

In order to evaluate the reproducibility of the NPQ score, we performed a Mann-Whitney test on each couple of scores from two consecutive repeats on the monoclonal populations used in the experiments presented in Figure 3 of the Main Text. As displayed in Figure S15, the hypothesis of equality of the underlying distributions of the data was rejected for the q̃I scores of ***Pop_qT*** (p-value $10^{-6}$) and $wt4a^{-}$ (p-value $10^{-3}$). For all the other samples, we failed to reject the equality hypothesis (p-values $> 0.04$). Therefore, we could use these experiments to study q̃E and q̃T by identifying the repeats as identical.

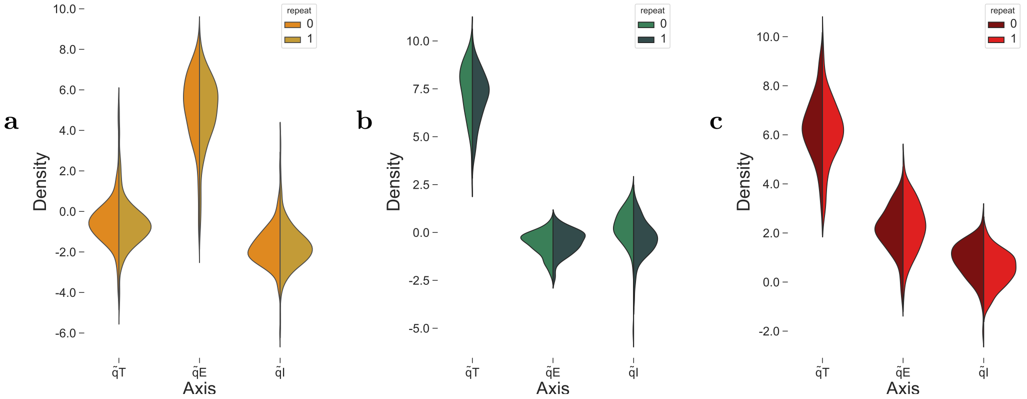

**Figure S15: Distribution of the NPQ scores for the two consecutive repeats (third and fourth) of the experiments on monoclonal populations. a**: ***Pop_qE*** monoclonal, two repeats – failed to reject equality ; **b**: ***Pop_qT*** monoclonal, two repeats – failed to reject equality except for q̃I (p-value $10^{-6}$); **c**: $wt4a^{-}$ HL-activated, monoclonal, two repeats – failed to reject equality except for q̃I (p-value $10^{-3}$).

#### 7.2 Main text figures

The various statistical tests performed on the elements presented in the Main Text are presented in Table S3. The null-hypothesis is described along with the p-values and the conclusion. We performed normality tests on the data and report on the sample sizes. The data compared often share the same size, except when comparing $D_{i,i}$ and $D_{i,j}$. In the latter case, we used a bootstraping method, by performing the test on subsets of $D_{i,j}$ with sizes equal to $D_{i,i}$ (100 draws). The tests were performed in Python with the SciPy statistics library functions [17].

| Figure | Population | Variable | Size | Null hypothesis | Samples compared | Test | p-value | Conclusion |
|---|---|---|---|---|---|---|---|---|
| 3B | ***Pop_qE*** monoclonal | $D_{i,i}$ | 176 | the central tendancy of the distribution of X is greater then Y's. | X = $D_{i,i}$ and Y = $D_{i,j}$ | Mann-Whitney bootstrap $D_{i,j}$ | $10^{-33}$ | rejected |
| | | $D_{i,j}$ | 30976 | | | | | |
| 3E | ***Pop_qT*** monoclonal | $D_{i,i}$ | 77 | the central tendancy of the distribution of X is greater then Y's. | X = $D_{i,i}$ and Y = $D_{i,j}$ | Mann-Whitney bootstrap $D_{i,j}$ | $10^{-9}$ | rejected |
| | | $D_{i,j}$ | 5929 | | | | | |
| 3A | repeat 0 | q̃E | 176 | the underlying distributions of q̃E in the two repeats are equal | two consecutive repeats | Mann-Whitney | 0.25 | failed to reject |
| | repeat 1 | q̃E | 176 | | | | | |
| 3E | repeat 0 | q̃T | 77 | the underlying distributions of q̃T in the two repeats are equal | two consecutive repeats | Mann-Whitney | 0.04 | failed to reject |
| | repeat 1 | q̃T | 77 | | | | | |
| 3C | ***Pop_qE*** | q̃E | | all input samples are from populations with equal variances | 3 non-monoclonal<br>2 monoclonal | Levene | 0.99 | failed to reject |
| 3F | ***Pop_qT*** | q̃T | | all input samples are from populations with equal variances | 4 non-monoclonal<br>1 monoclonal | Levene | $10^{-9}$ | rejected[1] |
| 3I | $wt4a^-$ monoclonal | $D_{i,i}$ | 77 | the central tendancy of the distribution of X is greater then Y's. | X = $D_{i,i}$ and Y = $D_{i,j}$ | Mann-Whitney bootstrap $D_{i,j}$ | $10^{-6}$ | rejected |
| | | $D_{i,j}$ | 5929 | | | | | |
| 3H | repeat 0 | q̃T | 77 | the underlying distributions of q̃T in the two repeats are equal | two consecutive repeats | Mann-Whitney | 0.4 | failed to reject |
| | repeat 1 | q̃T | 77 | | | | | |
| 3G | repeat 0 | q̃E | 77 | the underlying distributions of q̃E in the two repeats are equal | two consecutive repeats | Mann-Whitney | 0.5 | failed to reject |
| | repeat 1 | q̃E | 77 | | | | | |
| 4A | ***Pop_qT*** to $wt4a^-$ | q̃E, q̃T | 200-800 | the underlying distributions of q̃E and q̃T are uncorrelated | q̃E and q̃T for the total population | Pearson | $<10^{-250}$ | rejected |
| | | q̃E, q̃T | 200-800 | | q̃E and q̃T for individual sequentially activated populations | | $<10^{-37}$ | rejected |
| 4D | cc124 to *cc124 | q̃E, q̃T | 300-800 | the underlying distributions of q̃E and q̃T are uncorrelated | q̃E and q̃T for the total population | Pearson | $<10^{-230}$ | rejected |
| | | q̃E, q̃T | 300-800 | | q̃E and q̃T for individual sequentially activated populations | | <0.3 | failed to reject[2] |

**Table S3: Statistical tests performed on the elements presented in the Main Text.** [1] the test was rejected due to the diversity of variance in the non-monoclonal strains, but the monoclonal variance is not an outlier. [2] Accepted only once for 80 min of activation, with other values smaller than $10^{-40}$.

### 8 Evaluation of the relevance of the axis q̃I

We did not quantitatively analyze the slower NPQ component in the Main Text since this biological phenomenon is the hardest to control among the ones investigated and since no mutant strain exists. Accordingly, once we obtained the machine learning framework, we focused on populations where qI could be neglected.

Figure S16 visually demonstrates that the amplitude of characteristic feature for the qI, ie. global decrease of the fluorescence throughout the whole experiment with a final fluorescence level $F'_m$ not recovering to the initial value, increases as the corresponding datapoint moves away from the origin. Notably, on the q̃I axis, the distance of ***Pop_qI*** from the origin is only slightly larger than that of the other training populations, indicating that this NPQ component is smaller and not as tightly controlled as the other two in the reference dataset. This might reflect the photoinhibited state of *stt7-1* even under low-light growth conditions .

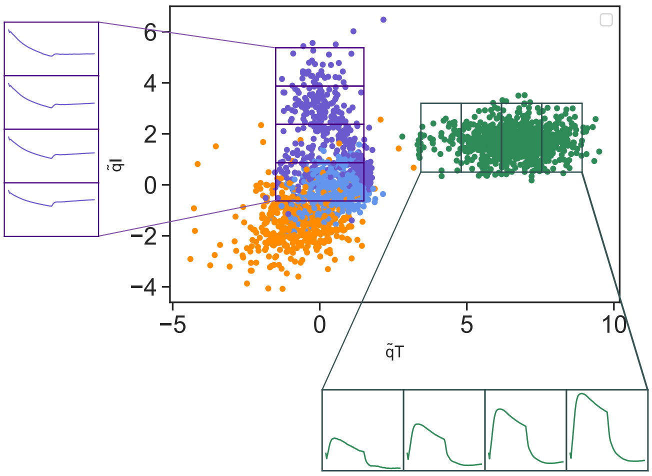

**Figure S16: Qualitative analysis of the evolution of the fluorescence traces along the q̃I and q̃T axis.** The purple population corresponds to the population ***Pop_qI***. The span of (98% of) the population along the q̃I axis is split into 4 equal parts, and the average fluorescence trace within each quarters are computed using the fluorescence traces (normalized to the first point) corresponding to the data points falling inside the quarter. The same is performed for the ***Pop_qT*** population (green point cloud) along the q̃T axis. For all plots, the span of the y-axis is equal to 0.7, with the first point falling at position 1.

#### References

[1] A. Lahlou et al. "Fluorescence to Measure Light Intensity". In: *Nature Methods* 20.12 (Dec. 2023), pp. 1930–1938. DOI: 10.1038/s41592-023-02063-y.

[2] R. Zhang et al. "Macroscale Fluorescence Imaging against Autofluorescence under Ambient Light". In: *Light: Science & Applications* 7.1 (Nov. 2018), p. 97. DOI: 10.1038/s41377-018-0098-6.

[3] J. Pech-Pacheco et al. "Diatom Autofocusing in Brightfield Microscopy: A Comparative Study". In: *Proceedings 15th International Conference on Pattern Recognition. ICPR-2000.* Vol. 3. Sept. 2000, 314–317 vol.3. DOI: 10.1109/ICPR.2000.903548.

[4] L. Girolomoni et al. "LHCSR3 Is a Nonphotochemical Quencher of Both Photosystems in Chlamydomonas Reinhardtii". In: *Proceedings of the National Academy of Sciences* 116.10 (Mar. 2019), pp. 4212–4217. DOI: 10.1073/pnas.1809812116.

[5] G. Peers et al. "An Ancient Light-Harvesting Protein Is Critical for the Regulation of Algal Photosynthesis". In: *Nature* 462.7272 (Nov. 2009), pp. 518–521. DOI: 10.1038/nature08587.

[6] K. Kosuge et al. "LHCSR1-dependent Fluorescence Quenching Is Mediated by Excitation Energy Transfer from LHCII to Photosystem I in Chlamydomonas Reinhardtii". In: *Proceedings of the National Academy of Sciences* 115.14 (Apr. 2018), pp. 3722–3727. DOI: 10.1073/pnas.1720574115.

[7] M. Cantrell, M. A. Ware, and G. Peers. "Characterizing Compensatory Mechanisms in the Absence of Photoprotective qE in Chlamydomonas Reinhardtii". In: *Photosynthesis Research* 158.1 (Oct. 2023), pp. 23–39. DOI: 10.1007/s11120-023-01037-7.

[8] E. Tyystjärvi and E.-M. Aro. "The Rate Constant of Photoinhibition, Measured in Lincomycin-Treated Leaves, Is Directly Proportional to Light Intensity". In: *Proceedings of the National Academy of Sciences of the United States of America* 93 (Apr. 1996), pp. 2213–8. DOI: 10.1073/pnas.93.5.2213.

[9] E. Tyystjärvi. "Chapter Seven - Photoinhibition of Photosystem II". In: *International Review of Cell and Molecular Biology.* Ed. by K. W. Jeon. Vol. 300. International Review of Cell and Molecular Biology. Academic Press, Jan. 2013, pp. 243–303. DOI: 10.1016/B978-0-12-405210-9.00007-2.

[10] M. Nilkens et al. "Identification of a Slowly Inducible Zeaxanthin-Dependent Component of Non-Photochemical Quenching of Chlorophyll Fluorescence Generated under Steady-State Conditions in Arabidopsis". In: *Biochimica et Biophysica Acta (BBA) - Bioenergetics* 1797.4 (Apr. 2010), pp. 466–475. DOI: 10.1016/j.bbabio.2010.01.001.

[11] R. Chouket et al. "Extra Kinetic Dimensions for Label Discrimination". In: *Nature communications* 13.1 (2022), pp. 1–8.

[12] Alienor134. *Alienor134/Image_segmentation:* Zenodo. Nov. 2021. DOI: 10.5281/zenodo.5684343.

[13] U. Schreiber. "Pulse-Amplitude-Modulation (PAM) Fluorometry and Saturation Pulse Method: An Overview". In: *Chlorophyll a Fluorescence: A Signature of Photosynthesis.* Ed. by G. C. Papageorgiou and Govindjee. Advances in Photosynthesis and Respiration. Dordrecht: Springer Netherlands, 2004, pp. 279–319. ISBN: 978-1-4020-3218-9. DOI: 10.1007/978-1-4020-3218-9_11.

[14] G. Allorent et al. "A Dual Strategy to Cope with High Light in Chlamydomonas Reinhardtii". In: *The Plant Cell* 25.2 (Feb. 2013), pp. 545–557. DOI: 10.1105/tpc.112.108274.

[15] M. Á. Ruiz-Sola and D. Petroutsos. "A Toolkit for the Characterization of the Photoprotective Capacity of Green Algae". In: *Plastids.* Ed. by E. Maréchal. Vol. 1829. New York, NY: Springer US, 2018, pp. 315–323. ISBN: 978-1-4939-8653-8 978-1-4939-8654-5. DOI: 10.1007/978-1-4939-8654-5_21.

[16] T. Roach and C. S. Na. "LHCSR3 Affects De-Coupling and Re-Coupling of LHCII to PSII during State Transitions in Chlamydomonas Reinhardtii". In: *Scientific Reports* 7.1 (Mar. 2017), p. 43145. DOI: 10.1038/srep43145.

[17] P. Virtanen et al. "SciPy 1.0: Fundamental Algorithms for Scientific Computing in Python". In: *Nature Methods* 17.3 (Mar. 2020), pp. 261–272. DOI: 10.1038/s41592-019-0686-2.
